# Mitochondrial fusion regulates proliferation and differentiation in the type II neuroblast lineage in *Drosophila*

**DOI:** 10.1101/2021.01.08.425832

**Authors:** Dnyanesh Dubal, Prachiti Moghe, Bhavin Uttekar, Richa Rikhy

**Affiliations:** Biology, Indian Institute of Science Education and Research, Homi Bhabha Road, Pashan, Pune, 411008, India; EMBL Heidelberg, Meyerhofstraße 1, 69117 Heidelberg, Germany

**Keywords:** mitochondria, *Drosophila*, neuroblast, differentiation, Opa1, Marf, Notch, Drp1

## Abstract

Optimal mitochondrial function determined by mitochondrial dynamics, morphology and activity is coupled to stem cell differentiation and organism development. However, the mechanisms of interaction of signaling pathways with mitochondrial morphology and activity are not completely understood. We assessed the role of mitochondrial fusion and fission in differentiation of neural stem cells called neuroblasts (NB) in the *Drosophila* brain. Depletion of mitochondrial inner membrane fusion protein Opa1 and mitochondrial outer membrane protein Marf in the *Drosophila* type II neuroblast lineage led to mitochondrial fragmentation and loss of activity. Opa1 and Marf depletion did not affect the numbers and polarity of type II neuroblasts but led to a decrease in proliferation and differentiation of cells in the lineage. On the contrary, loss of mitochondrial fission protein Drp1 led to mitochondrial fusion but did not show defects in proliferation and differentiation. Depletion of Drp1 along with Opa1 or Marf also led to mitochondrial fusion and suppressed fragmentation, loss of mitochondrial activity, proliferation and differentiation in the type II NB lineage. We found that Notch signaling depletion via the canonical pathway showed mitochondrial fragmentation and loss of differentiation similar to Opa1 mutants. An increase in Notch signaling required mitochondrial fusion for NB proliferation. Further, Drp1 mutants in combination with Notch depletion showed mitochondrial fusion and drove differentiation in the lineage suggesting that fused mitochondria can influence Notch signaling driven differentiation in the type II NB lineage. Our results implicate a crosstalk between Notch signalling, mitochondrial activity and mitochondrial fusion as an essential step in type II NB differentiation.

## Introduction

Mitochondria are sparse and fragmented in stem cells and occur as an elaborate network in differentiated cells [1–3]. Stem cells largely depend upon glycolysis as an energy source, whereas differentiated cells produce a large amount of ATP by electron transport chain (ETC) activity [1,4,5]. Fused mitochondrial morphology is associated with high membrane potential, increased ETC activity and high ATP production, while low membrane potential, reduced ETC activity and low ATP production is seen in fragmented mitochondria [1]. Mitochondrial architecture is regulated by a balance of fusion and fission events [6]. Proteins belonging to the family of large GTPases are involved in mitochondrial fusion and fission. Optic atrophy 1 (Opa1, or Opa1-like in *Drosophila*) and Mitofusin or Mitochondrial assembly regulatory factor (Marf in *Drosophila* or Mitofusin, Mfn in mammals) facilitate inner and outer mitochondrial membrane fusion respectively while Dynamin related protein 1 (Drp1) is required for mitochondrial fragmentation [7–10]. A balance of levels and activity of these proteins regulates mitochondrial shape in the cell [6]. Further, Opa1 plays a significant role in regulating cristae organization in addition to inner membrane fusion [11]. Opa1 oligomerization and inner membrane cristae organization is important for ETC activity. The presence of elaborate cristae leads to organization of ETC complexes as super complexes and enhances their activity [12]. Hyperfusion of mitochondria protects them from degradation in autophagy and also loss of ETC activity [13–16].

Recent studies show that alteration of mitochondrial dynamics affects signaling pathways such as the Notch signaling pathway during stem cell differentiation. The Notch receptor is a transmembrane protein activated by ligands such as Delta. The Delta-Notch interaction is followed by cleavage of the Notch intracellular domain (NICD) in the signal receiving cell. NICD enters the nucleus and regulates gene expression along with Suppressor of hairless (Su(H)) by the canonical pathway thereby providing a signal for proliferation or differentiation [17]. Fragmented mitochondrial morphology maintained by Drp1 in ovarian follicle cells in *Drosophila* is crucial for activating Notch signaling [18,19]. Similarly, loss of Opa1 and Mfn leading to mitochondrial fragmentation in mouse embryonic stem cells causes hyperactivation of Notch and reduces differentiation of ESCs into functional cardiomyocytes due to loss of calcium buffering [20]. On the other hand, activation of Notch signaling by depletion of mitochondrial fusion and increasing reactive oxygen species (ROS) enhances differentiation in mammalian neural stem cells [21]. Thus mitochondrial fragmentation along with elevated calcium and reactive oxygen species increase has been found to be involved in Notch signaling in these contexts. It is of interest to understand whether Notch signaling induces appropriate mitochondrial morphology in differentiation.

The *Drosophila* neural stem cell or neuroblast (NB) differentiation model has been used effectively to identify regulators of steps of differentiation such as stem cell renewal, asymmetric cell division, polarity formation and lineage development. NBs rely on glycolysis and ETC activity for their energy production during differentiation and tumorigenesis [22–24]. Mitochondrial fusion has recently been found to be essential for tumorigenesis [24]. It remains to be studied whether mitochondrial morphology is also regulated to provide appropriate activity for differentiation in NBs. The type I NB lineage in the larval brain consists of 90 NBs marked by the expression of the transcription factor Deadpan (Dpn). These NBs divide asymmetrically to give rise to a ganglion mother cells (GMCs) marked by the expression of Prospero (Pros) in the nucleus [25,26]. NBs of the type II lineage also express Dpn and are 8 in number [27]. Like the mammalian neural stem cell differentiation type II NBs undergo multiple steps of differentiation by forming transit amplifying cells called intermediate neural precursor cells (INPs). Newly formed INPs are smaller in size as compared to type II NBs and do not express Dpn. INPs undergo a defined series of transcriptional changes to form mature INPs (mINPs). mINPs express Dpn and proliferate to form GMCs that express Pros. GMCs in both the type I and type II lineages finally differentiate into neurons or glia (Figure 1A). Notch signaling regulates type II NB number and differentiation in the type II NB lineage [28,29].

**Figure 1:**
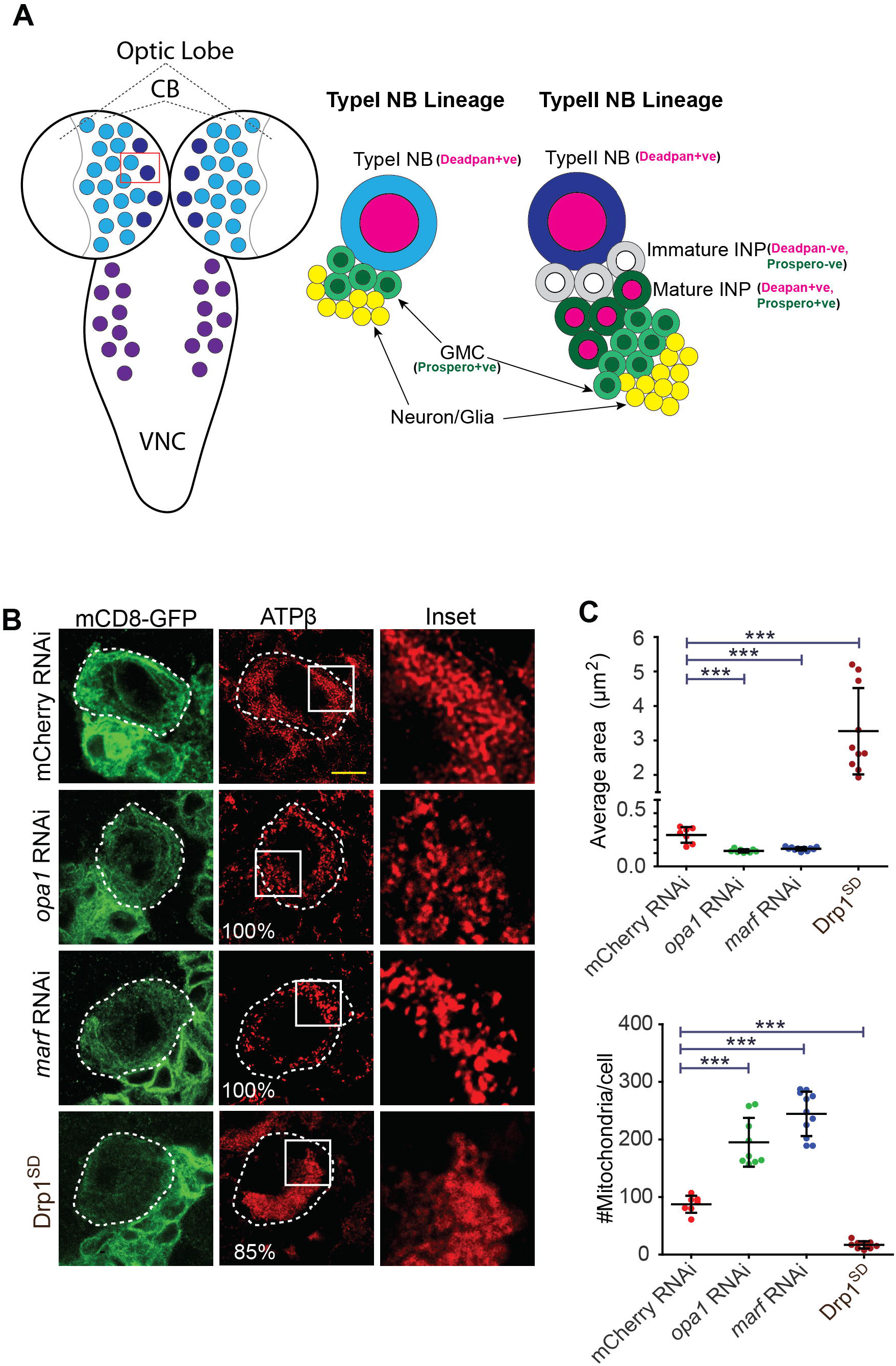
Mitochondrial morphology protein depletion leads to altered mitochondrial distribution in neuroblasts. A: Schematic of larval CNS (left) containing the central brain (CB) lobes and ventral nerve cord (VNC) and type I (blue) and type II NB (purple) distribution and lineages of type I (blue, right) and type II NB (purple, right). The type I NB lineage has Dpn (magenta nuclei) positive NBs and Pros (green nuclei) positive GMCs. The type II NB lineage has Dpn positive NBs (magenta nuclei), Dpn negative and Pros negative immature INPs (black and white), Dpn positive mINPs (magenta nuclei) and Pros positive GMCs (green nuclei). Dpn positive NBs are clearly distinguishable as larger cells as compared to Dpn positive mINPs in each type II NB lineage. B-D: Mitochondrial morphology and distribution in type II NBs (white dotted line, magnified area shown in the panel on the right) stained with ATPβ (red) antibody using STED super resolution microscopy is shown in representative images with zoomed inset in the right panel (B). mCherry RNAi (100% tubular, 32 NBs, 8 brains), *opa1* RNAi (100% fragmented, 46,8), *marf* RNAi (100% fragmented, 45,8), Drp1^SD^ (85% clustered, 80,10). Average mitochondrial area quantification from type II NBs (C) in mCherry RNAi (7 type II NBs, 3 brains), *opa1* RNAi (9,3), *marf* RNAi (11,3), Drp1^SD^(10,3). Mitochondrial number quantification in mCherry RNAi (7,3), *opa1* RNAi (9,3), *marf* RNAi (11,3), Drp1^*SD*^ (10,3) (D). Scale bar-5μm C-D: Graphs show mean + sd. Statistical analysis is done using an unpaired t-test. **-p<0.01, ***-p<0.001

In this study we have assessed the role of mitochondrial morphology proteins Opa1, Marf and Drp1 in regulating type II NB differentiation. We find that Opa1 and Marf mediated mitochondrial fusion is essential for type II NB differentiation. RNAi mediated knockdown of mitochondrial fusion proteins Opa1 and Marf led to mitochondrial fragmentation, loss of mitochondrial activity and defects in differentiation in the type II NB lineage while NB number and polarity remained unaffected. On the other hand there was no defect in differentiation in NBs depleted of mitochondrial fission protein Drp1. Inhibition of mitochondrial fragmentation in Opa1 and Marf mutants by additional depletion of Drp1 suppressed the differentiation defects suggesting that fused mitochondria are essential for type II NB differentiation. Further, Notch depletion led to fragmented mitochondria and loss of differentiation. Increased Notch activity showed mitochondrial fusion and mitochondrial fusion in the type II NB lineage deficient of Notch led to differentiation. Our results show that mitochondrial fusion interacts with Notch signaling to drive differentiation in the type II NB lineage.

## Results

### Depletion of Opa1 and Marf leads to mitochondrial fragmentation and depletion of Drp1 leads to mitochondrial fusion in type II NBs

We depleted Opa1, Marf and Drp1 to investigate the effect of perturbation of mitochondrial morphology on NB numbers and differentiation. We expressed multiple RNAi lines and mutants against *opa1*, *marf* and *drp1* with neuronal Gal4 drivers in different stages of NB differentiation to analyze their effect on lethality and behavior (Figure S1). *inscuteable*-Gal4, *worniu*-Gal4, *scabrous*-Gal4 and *prospero*-Gal4 were used to deplete Opa1, Marf and Drp1 using RNAi expression and Drp1 using a dominant negative mutant in all NBs. Opa1, Marf and Drp1 depletion by multiple RNAi lines and mutants showed survival of animals until the pupal stage with *inscuteable*-Gal4 and *worniu*-Gal4 and were lethal or showed behavioral defects as adults. The RNAi lines for Opa1 and Marf that gave a stronger defect with *inscuteable*-Gal4 and *worniu*-Gal4 and the dominant negative mutant of Drp1, (Drp1^SD^) [30] were used to deplete these proteins in the type II NB lineage using *pointed*-Gal4 for further experiments (Figure 1A, S1). *opa1* RNAi, *marf* RNAi and Drp1^SD^ expression in the type II NB lineage gave normal adults at 25 ^o^C. *opa1* RNAi and *marf* RNAi expression in type II NB lineage when performed at a higher temperature of 29°C with *pointed*-Gal4 gave sluggish adults.

Mitochondria are tubular and uniformly distributed around the nucleus in type I NBs [31]. To characterize mitochondrial morphology in type II NBs, we performed super-resolution Stimulated Emission Depletion microscopy (STED) on mitochondria stained with antibody against ATPβ subunit of complex V in the third instar larval brain. We observed thread-like mitochondria evenly distributed around the nucleus in control type II NBs (Figure 1B, S2A). We used *pointed*-Gal4, mCD8-GFP to identify the type II NBs using mCD8-GFP and deplete Opa1, Marf and Drp1. RNAi against mitochondrial fusion proteins Opa1 and Marf has been previously shown to deplete the corresponding mRNA and lead to mitochondrial fragmentation in electron microscopy studies [32–35]. We observed a distinct shift in mitochondrial morphology from a tubular organization to dispersed punctae on depletion of Opa1 and Marf using two different RNAi lines as compared to *mcherry*-RNAi controls, confirming the requirement of these proteins for mitochondrial fusion (Figure 1B, S2A). We quantified the numbers and area of optically resolvable mitochondria in STED images to estimate the mitochondrial size in each NB. NBs depleted of Opa1 and Marf showed a significant increase in mitochondrial numbers and decrease in mitochondrial area as compared to controls (Figure 1C-D, S2B-C). Interestingly, the extent of mitochondrial fragmentation was similar upon depletion of either Opa1 or Marf. Further, expression of Drp1^SD^ resulted in clustering of mitochondria on one side of the NB suggesting that mitochondria were fused (Figure 1B). Drp1 depletion led to a significant increase in mitochondrial area and decrease in mitochondrial numbers as compared to controls (Figure 1C-D). In summary, depletion of Opa1 and Marf led to mitochondrial fragmentation and depletion of Drp1 led mitochondrial fusion in the type II NBs.

### Mitochondrial fusion is required for differentiation in the type II NB lineage

We checked the effect of knockdown of mitochondrial morphology proteins on NB number, polarity and differentiation. We depleted mitochondrial morphology proteins using *worniu*-Gal4 to estimate the numbers of type I and II NBs. The overall number of NBs was not changed upon depletion of Opa1, Marf and Drp1 (Figure S3A-B) and the type II number also remained unaffected (Figure S3C). Apico-basal distribution of polarity proteins Bazooka and Numb in metaphase is essential for fate determination in the lineage [26]. We found that the NBs did not show a defect in polarised distribution of Bazooka and Numb on depletion of Opa1, Marf or Drp1 (Figure S3D).

We identified cells in each type II NB lineage based on their specific molecular profile (Figure 1A). Dpn immunostaining was used to count mINPs (Figure 2A) while nuclear Pros immunostaining was used to count GMCs (Figure 2B) in type II NB lineage. Immature INPs lacked both Dpn and Pros. Immature INPs were similar in number in Opa1, Marf and Drp1 depleted type II NB lineages (Figure 2C). On the other hand, mINPs and GMCs were lowered on expression of two different RNAi lines against *opa1* as compared to controls (Figure 2A,B,D,E, S2D-G). Marf depletion by different RNAi lines did not alter mINP numbers but reduced GMCs as compared to controls (Figure 2A,B,D,E, S2D-G). There was no change in mINPs and GMCs in each lineage in Drp1^SD^ expressing type II NBs (Figure 2A,B,D,E). The numbers of GMCs were also reduced in the *opa1* RNAi but not the *marf* depleted type I NB lineages (Figure S3E-F). Loss of mitochondrial fusion in Opa1 and Marf mutants may give rise to defective INPs thereby impairing differentiation in the type II NB lineage.

**Figure 2:**
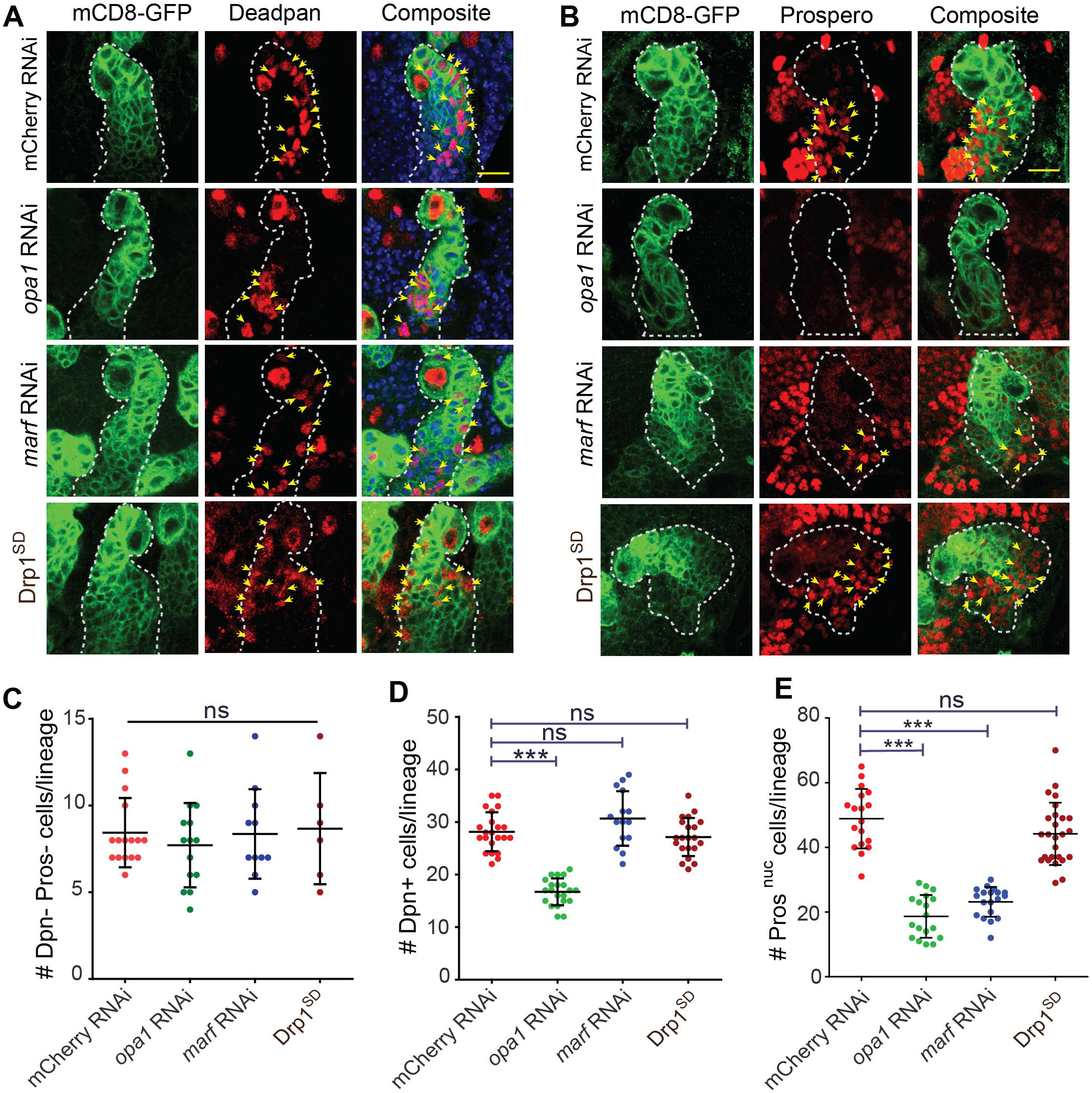
Depletion of Opa1 causes loss of differentiated mINPs and GMCs in type II NB lineage. A-B: Representative confocal images of the type II NB lineage showing that Dpn positive mINPs (red, yellow arrows) are reduced in *pnt*-Gal4, UAS-mCD8-GFP driven *opa1* RNAi (A) and Pros (red, yellow arrows) positive GMCs (B) are lowered in *opa1* RNAi and *marf* RNAi expressing type II NBs C: Quantification of immature INPs (Dpn and Pros negative) per type II NB lineage. mCherry RNAi (8 type II NB lineages, 4 Brains), *opa1* RNAi (14,5), *marf* RNAi (11,3), Drp1^SD^ (6,3). D: Quantification of Dpn positive mINPs (yellow arrows) per type II NB lineage in mCherry RNAi (n=22 type II NB lineages, 4 brains), *opa1* RNAi (21,6), *marf* RNAi (13,8), Drp1^SD^ (28,10). E: Quantification of Pros positive GMCs (yellow arrows point to nuclear Pros) per type II NB lineage (E) of mCherry RNAi (18 type II NB lineages,5 brains), *opa1* RNAi (18,5), *marf* RNAi (19,5), Drp1^SD^ (26,9). Scale bar-10μm C, D, E: Graphs show mean + sd. Statistical analysis is done using an unpaired t-test. ns- non-significant, **-p<0.01, ***-p<0.001.

In summary, depletion of Opa1 decreased differentiation in the type II NB lineage to a greater extent compared to Marf, despite comparable disruption of mitochondrial morphology to fragmentation. Opa1 and Marf depletion may give rise to loss of cells in the lineage due to defects in the type II NB and mINP division. Depletion of mitochondrial fission protein Drp1 did not show any defects in differentiation. The phenotype of increased mitochondrial fragmentation and delay in NB division thereby leading to loss of differentiation has been previously noted on depletion of ETC components [23]. Apart from mitochondrial fusion, Opa1 is also involved in maintenance of cristae architecture [11,36–38]. It is therefore possible that a specific defect in mitochondrial inner membrane organization caused by Opa1 depletion in addition to loss of fusion is a cause for a loss in mINPs and GMCs in the type II NB lineage.

### Depletion of mitochondrial fission protein Drp1 along with Opa1 or Marf shows fused mitochondria and suppresses the defects in differentiation

Mitochondrial fragmentation in *opa1* and *marf* depleted cells requires the activity of mitochondrial fission protein Drp1 [30,32,39,40]. To alleviate the mitochondrial fragmentation defect seen in *opa1* and *marf* knockdown, we made combinations of Drp1^SD^;mRFP (data not shown), Drp1^SD^*;opa1* RNAi and Drp1^SD^*;marf* RNAi and analyzed mitochondrial morphology. Unlike *opa1* RNAi and *marf* RNAi (Figure 1), the Drp1^SD^*;opa1* RNAi and Drp1^SD^*;marf* RNAi combinations showed clustered mitochondrial morphology. The mitochondrial cluster was more resolved as compared to Drp1^SD^ alone and Drp1^SD^, mRFP (data not shown), with a few punctate mitochondria appearing and a small but significant decrease in average mitochondrial area in double mutants (Figure 3A-C).

**Figure 3:**
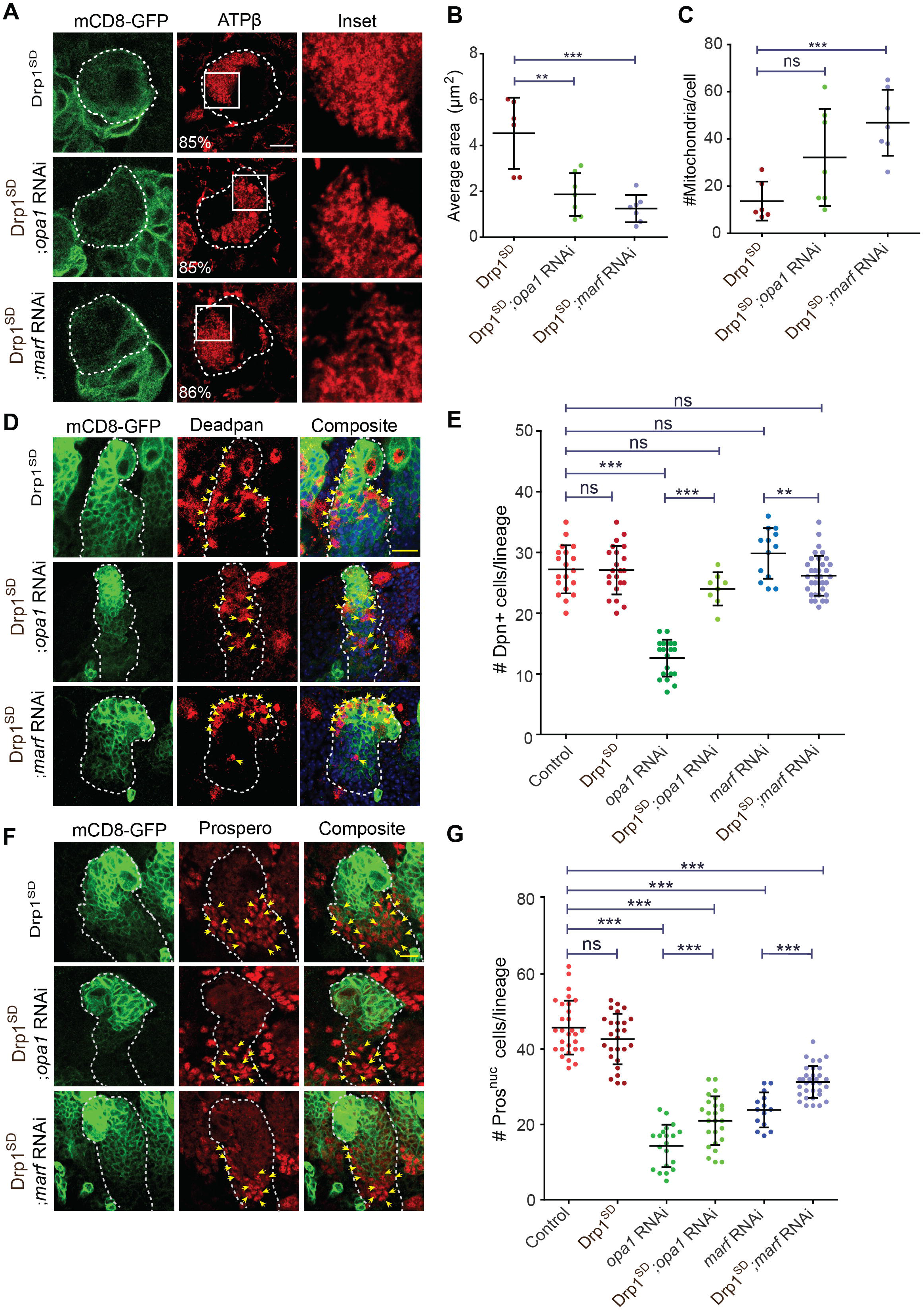
Drp1 depletion in *opa1* RNAi and *marf* RNAi expressing type II NB lineages leads to a suppression of the differentiation defect. A-C: Representative images with zoomed inset in the right panel of type II NBs (white dotted line) containing *pnt*-Gal4, mCD8-GFP (green) stained for mitochondrial morphology with ATPβ (red) and imaged using STED (A). *pnt*-Gal4, UAS-mCD8-GFP with Drp1^SD^ (85% clustered, 80 NBs,10 Brains), Drp1^SD^;*opa1* RNAi (85% clustered, 96,12), Drp1^SD^;*marf* RNAi (86% clustered, 96,12). Quantification of average mitochondrial area (B) in type II NB of Drp1^*SD*^ (6,3), Drp1^SD^;*opa1* RNAi (7,4), Drp1^SD^;*marf* RNAi (7,4). Quantification of mitochondrial numbers (C) in type II NB of Drp1^SD^ (6,3), Drp1^SD^;*opa1* RNAi (7,4), Drp1^SD^;*marf* RNAi (7,4). Scale bar-10μm. D-E: Type II NB lineages (yellow dotted line) showing expression of mCD8-GFP (green) and Dpn (red, yellow arrows) (D). Quantification of Dpn positive mINPs (E) in control (23 NB lineages,16 brains), Drp1^SD^ (28,10), *opa1* RNAi (20,8), Drp1^SD^;*opa1* RNAi (8,6), *marf* RNAi (13,8), Drp1^SD^;*marf* RNAi (35,8). Scale bar-10μm. F-G: Type II NBs lineages (yellow dotted line) showing expression of mCD8-GFP (green) and Pros (red, yellow arrows point to nuclear Pros) (F). Quantification of Dpn positive mINPs (G) in control (29 NB lineages,18 brains), Drp1^SD^ (26,8), *opa1* RNAi (20,8), Drp1^SD^;*opa1* RNAi (26,8), *marf* RNAi (14,8), Drp1^SD^;*marf* RNAi (33,8). Scale bar-10μm B,C,E & G: Graphs show mean + sd. Comparative analysis was done by using unpaired t-test. ns-non significant, **-p<0.01, ***-p<0.001

We analyzed these double mutants for differentiation of type II NB lineage by staining larval brains with Dpn and Pros. The numbers of Dpn positive mINPs increased in Drp1^SD^*;opa1* RNAi as compared to *opa1* alone and were not significantly different from controls (Figure 3D-E). mINPs were similar to controls in *marf* RNAi and Drp1^SD^*;marf* RNAi mutants (Figure 3D-E). The Drp1^SD^*;opa1* RNAi combination showed an increase in Pros positive GMCs but did not completely restore their numbers (Figure 3F-G). GMC numbers also increased in the Drp1^SD^*;marf*^i^ combination as compared to *marf* RNAi (Figure 3F-G). In summary, the differentiation defect observed in *opa1* and *marf* depletion was partially reversed on mitochondrial fusion by additional depletion of Drp1.

### Decrease in mitochondrial activity in Opa1 and Marf depleted type II NBs is suppressed by additional depletion of Drp1

Change in mitochondrial morphology can adversely affect mitochondrial membrane potential and activity. Mitochondrial membrane potential (MMP) is a readout for mitochondrial quality and functionality [41]. We checked the effect of mitochondrial dynamics proteins depletion on MMP by using the potentiometric dye, Tetra-methyl-rhodamine methyl ester (TMRM) in *vivo* in living larval brains by live imaging. We estimated the relative fluorescence obtained from uptake of TMRM in mCD8-GFP marked type II NBs as compared to neighboring unmarked controls. We found a significant decrease in TMRM fluorescence and therefore MMP on depletion of Opa1 and Marf whereas in Drp1^SD^ expression did not affect the MMP (Figure 4A-B). Co-depletion of Drp1 along with Opa1 or Marf RNAi suppressed the MMP defect showing that outer mitochondrial membrane fusion restores mitochondrial membrane potential (Figure 4A-B).

**Figure 4:**
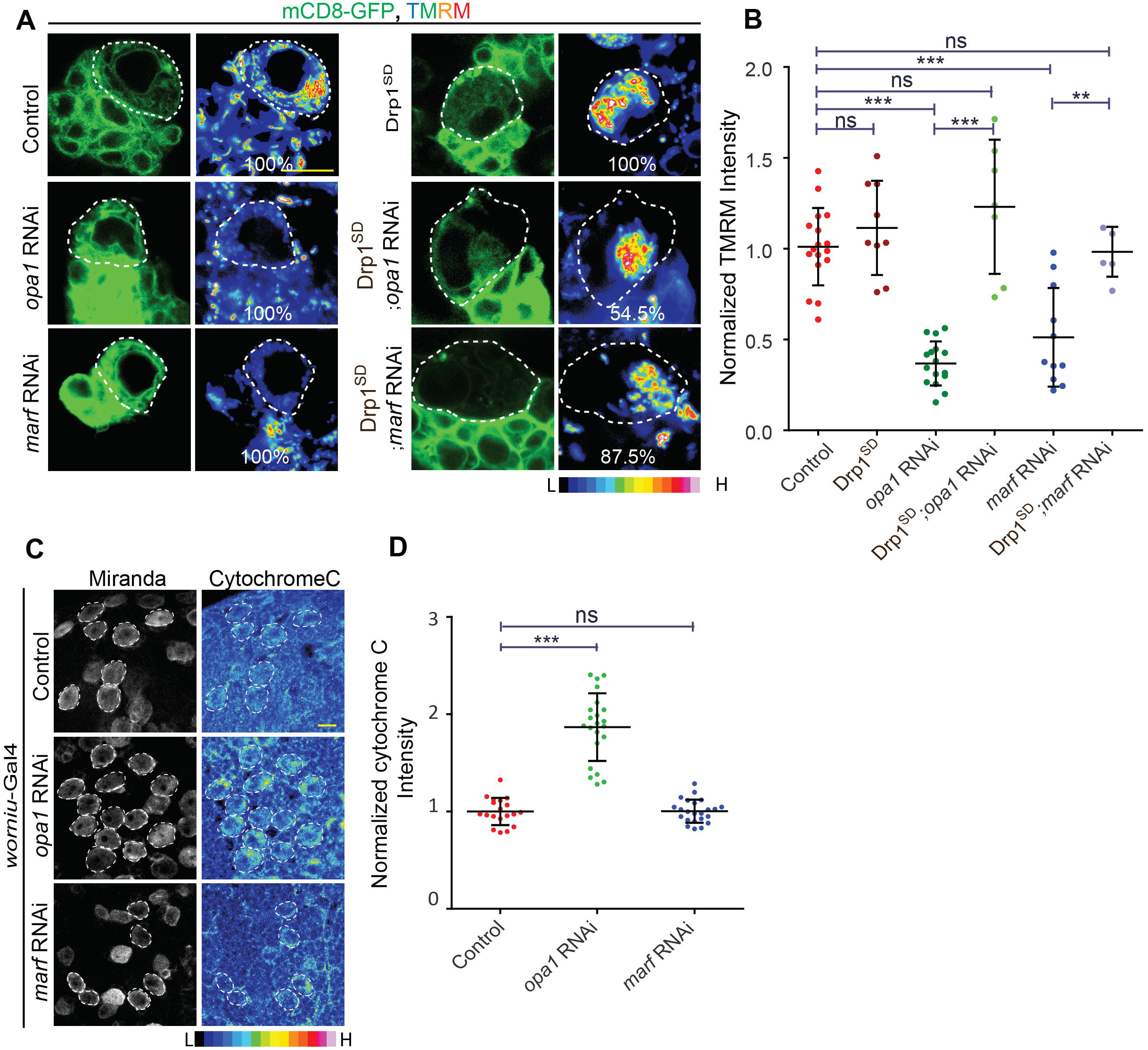
Co-depletion of Drp1 in Opa1 or Marf depleted type II NBs rescues the reduced mitochondrial membrane potential. A-B: Representative images showing decreased TMRM intensity in *opa1* RNAi and *marf* RNAi expressing type II NBs. *pnt*-Gal4, UAS-mCD8-GFP control (17 type II NBs, 8 brains), Drp1^SD^ (9,5), *opa1* RNAi (16,5), Drp1^SD^;*opa1* RNAi (11,4), *marf* RNAi (11,5), Drp1^SD^;*marf* RNAi (7,4). Scale bar-10μm (A). Graph showing normalized TMRM intensities in control (17 type II NBs, 8 brains), Drp1^SD^ (9,5), *opa1* RNAi (16,5), Drp1^SD^;*opa1* RNAi 7,4), *marf* RNAi (11,5), Drp1^SD^;*marf* RNAi (6,4) (B). C-D: Representative confocal images of larval NBs (C) showing increased cytochrome C staining in *opa1* RNAi using *worniu*-Gal4. Analysis of cytochrome C intensity normalised with control (D). Control (8 brains), *opa1* RNAi (7 brains), *marf* RNAi (7 brains). B,D: Graphs show mean + sd. Comparative analysis was done by using unpaired t-test. ns-non significant, **-p<0.01, ***-p<0.001

Decrease in MMP observed in Opa1 and Marf may result in drop in ATP levels in the cell that could trigger a stress response. We have previously found that change in ETC activity but not mitochondrial fusion obtained by Drp1 depletion causes ATP stress in *Drosophila* embryos [30,42]. Depletion of ATP causes an increase in AMP levels which further triggers phosphorylation of AMP activated protein kinase (pAMPK). pAMPK levels elevate in energy deprived conditions and act as an energy sensor inside the cell [43]. To check whether ATP stress was seen in mitochondrial fusion mutant NBs, we stained brains with antibodies against pAMPK. NB differentiation relies at least in part on glycolytic metabolism for ATP and loss of ATP in ETC mutants is seen when depleted of both glycolysis and ETC activity [22,23]. Therefore as a positive control for reduction in glycolysis and induction of pAMPK we added 2-deoxy-glucose (2-DG), a non-hydrolysable analogue of glucose to larval brains after dissection and observed a significant increase in pAMPK levels throughout the brain. However, we did not observe a change in pAMPK levels in *opa1* and *marf* mutant type II NBs as compared to neighboring NBs indicating that ATP stress similar to 2-DG treatment was not seen in these mutants (Figure S4A-B). This shows that depletion of ATP as read out by decrease in mitochondrial membrane potential in NBs was much lower as compared to that seen when glycolysis was inhibited with 2-DG.

Another consequence of change in mitochondrial morphology is alteration in the levels of reactive oxygen species (ROS) [44,45]. We estimated ROS levels using Dihydroxy ethidium (DHE) fluorescence in NBs as compared to neighboring cells. Consistent with previous studies [34,46], we observed increased DHE fluorescence indicating increased ROS on *opa1* knockdown and Drp1^SD^ expression (Figure S4C-D). *marf* depletion however did not change the DHE fluorescence significantly (Figure S4C-D). Since increased ROS was found in both *opa1* and *drp1* mutant NBs, and there was no effect on differentiation on Drp1^SD^ expression (Figure 2), we conclude that ROS is unlikely to play a significant role in differentiation in type II NBs. We further checked if a general increase in ROS can affect development. We found that in type I and II NBs expressing a mutant of human Superoxide dismutase (hSOD) [30,47], ROS was increased significantly (Figure S4C-D). However, hSOD expression in all NBs did not affect viability of flies and adults emerged similar to controls leading us to conclude that it did not impact differentiation and functionality of neurons (data not shown).

Even though both Opa1 and Marf depletion led to an equivalent fragmentation and loss of MMP of mitochondria in NBs, Opa1 depletion showed a specific decrease in mINPs as compared to Marf knockdown. Opa1 oligomerization leads to stabilization of cristae and localization of cytochrome c in a packed manner in cristae. Increase in cytochrome c occurs when Opa1 oligomerization is decreased and this correlates with loose cristae organization [36]. We visualized the distribution of cytochrome c in *opa1* and *marf* mutant NBs. Increase in cytochrome c was seen in *opa1* RNAi as compared to *marf* RNAi and controls (Figure 4C-D). This suggests that loosening of cristae architecture potentially causes specific spread of the cytochrome c signal on Opa1 depletion. Altogether, loss of MMP and increased cytochrome c in type II NBs deficient of *opa1* is suggestive of disruption of inner mitochondrial membrane architecture and activity in addition to fusion leading to loss of differentiation.

### Proliferation defects in Opa1 depleted type II NBs are suppressed by mitochondrial fusion on additional depletion of Drp1

The decreased lineage size observed in *opa1* and *marf* mutant type II NBs may result from apoptosis or lowered proliferation rates of each NB thereby reducing the numbers of INPs and GMCs in each lineage. Increased cytochrome C and ROS in *opa1* depletion suggests that type II NB differentiation could be decreased by apoptosis in the lineage [45,48–50]. For probing apoptosis, we assessed the levels of cleaved caspase in *opa1* mutants. Cleaved caspase staining was not elevated in *opa1* mutant lineages and was similar to controls (Figure S4E). Selectively marking apoptotic nuclei with the Terminal deoxynucleotidyl transferase (TdT) dUTP Nick-End Labeling (TUNEL) assay did not show any significant difference between Opa1 mutant NBs and control. We validated this assay by using UAS-hid expression as a positive control for apoptosis induction (Figure S4F-G). Therefore, cell death is not responsible for the differentiation defect in *opa1* mutant type II NBs.

mINPs divide asymmetrically to produce GMCs in the type II NB lineage. We analyzed the numbers of cells in mitosis in each type II NB lineage by staining for phospho-histone 3 (pH3) antibody. We found a significant reduction of pH3 positive cells in the type II lineage in *opa1* mutants while *marf* and *drp1* depletion did not show any significant change as compared to controls (Figure 5A-B). The numbers of pH3 positive cells were increased in the Drp1^SD^*;opa1* RNAi combination as compared to *opa1* RNAi and controls. The Drp1^SD^*;marf* RNAi combination also showed significant increase in pH3 positive cells per lineage as compared to controls indicating an increase in proliferation of mINPs in this combination (Figure 5A-B). The loss of pH3 positive cells showed that Opa1 depletion decreased the rate of mINP proliferation in the type II NB. This defect of loss of proliferation in Opa1 mutants was suppressed on outer mitochondrial fusion in Drp1 mutants.

**Figure 5:**
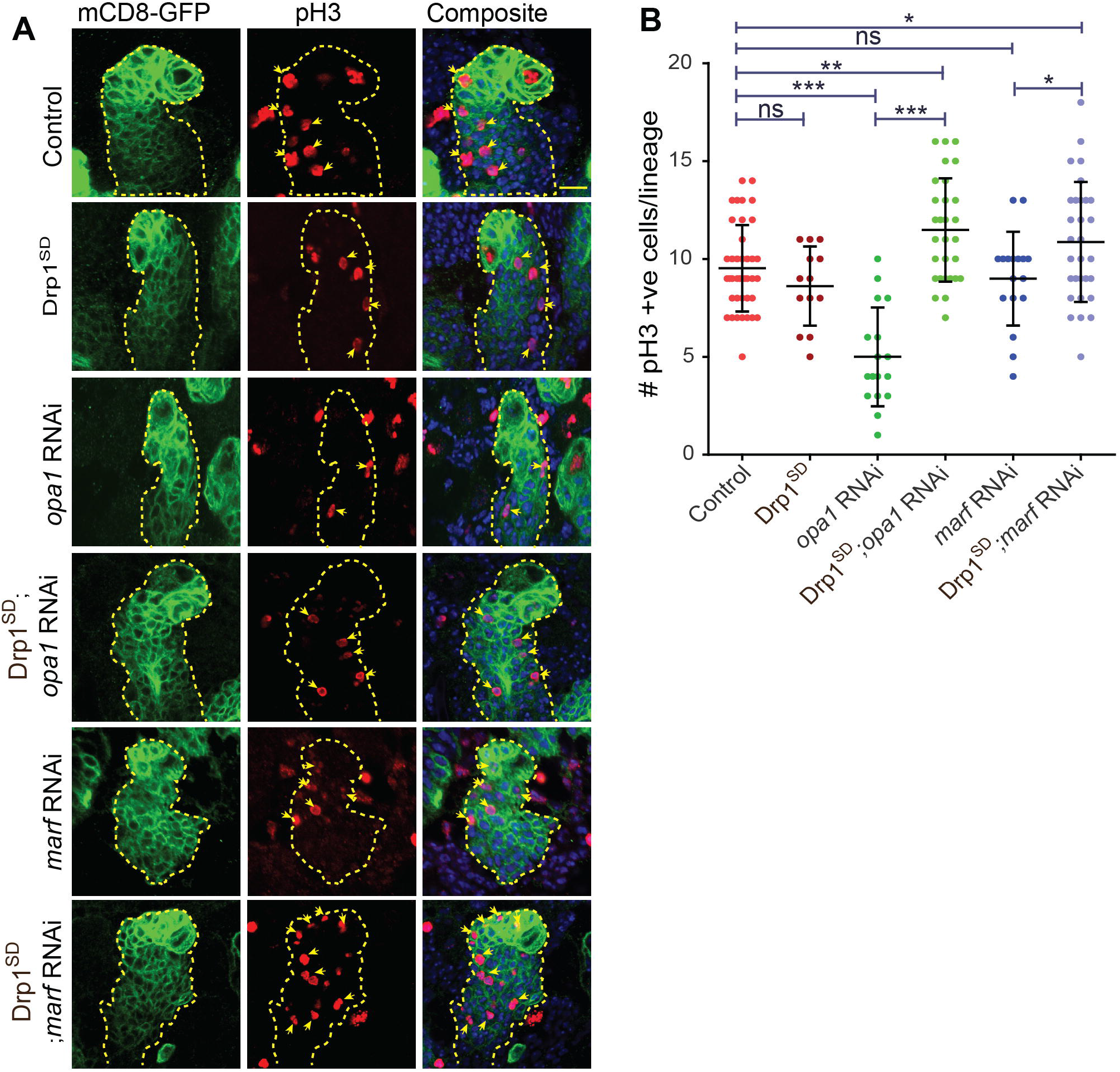
Proliferation defects in Opa1 depleted type II NBs are suppressed by inhibition of Drp1. A-B: Representative confocal images of type II NB lineages show loss of pH3 positive cells on *opa1* RNAi expressed with *pnt*-Gal4, UAS-mCD8-GFP in the type II NB lineage (A). Quantification of pH3 positive cells (B) in control (28 NB lineages,6 brains), Drp1^SD^ (13,3), *opa1* (17,3), Drp1^SD^;*opa1* RNAi (29,4), *marf* RNAi (17,3), Drp1^SD^;*marf* RNAi (30,7). Scale bar-10μm. B: Graph shows mean + sd. Statistical analysis was done by using unpaired t-test. ns-non significant, *-p<0.1, **-p<0.01, ***-p<0.001

### Notch signaling drives differentiation and fused mitochondrial morphology in the type II NB lineage

Notch signaling regulates numbers of NBs and their differentiation in the type II lineage in the *Drosophila* third instar larval brain [28,29,51]. Notch signaling is activated in type II NBs and suppressed in INPs. Notch signaling is again activated in the mINPs and GMCs in the lineage. Loss of Notch signaling leads to transformation of the type II NB to the type I NB with absence of Dpn positive mINPs. We overexpressed full length Notch (Notch^FL^) to increase Notch signaling and *notch* RNAi to decrease Notch signaling in the type II NB lineage. As documented previously [52], we observed an increased number of Dpn positive NBs on Notch^FL^ expression and a decreased number of type II NBs in *notch* RNAi expressing type II NBs. However, depletion of mitochondrial morphology proteins did not change the numbers of NBs (Figure S3A-C). Therefore, we conclude that the Notch signaling needed for maintenance of type II NB numbers was not defective in these mutants. We further analyzed the numbers of Dpn positive mINPs in lineages of remaining type II NBs that were present on *notch* RNAi expression. As expected, we observed decreased Dpn positive mINPs in *notch* RNAi while they remained unaffected in Notch^FL^ expressing type II NB lineage (Figure 6A-B).

**Figure 6:**
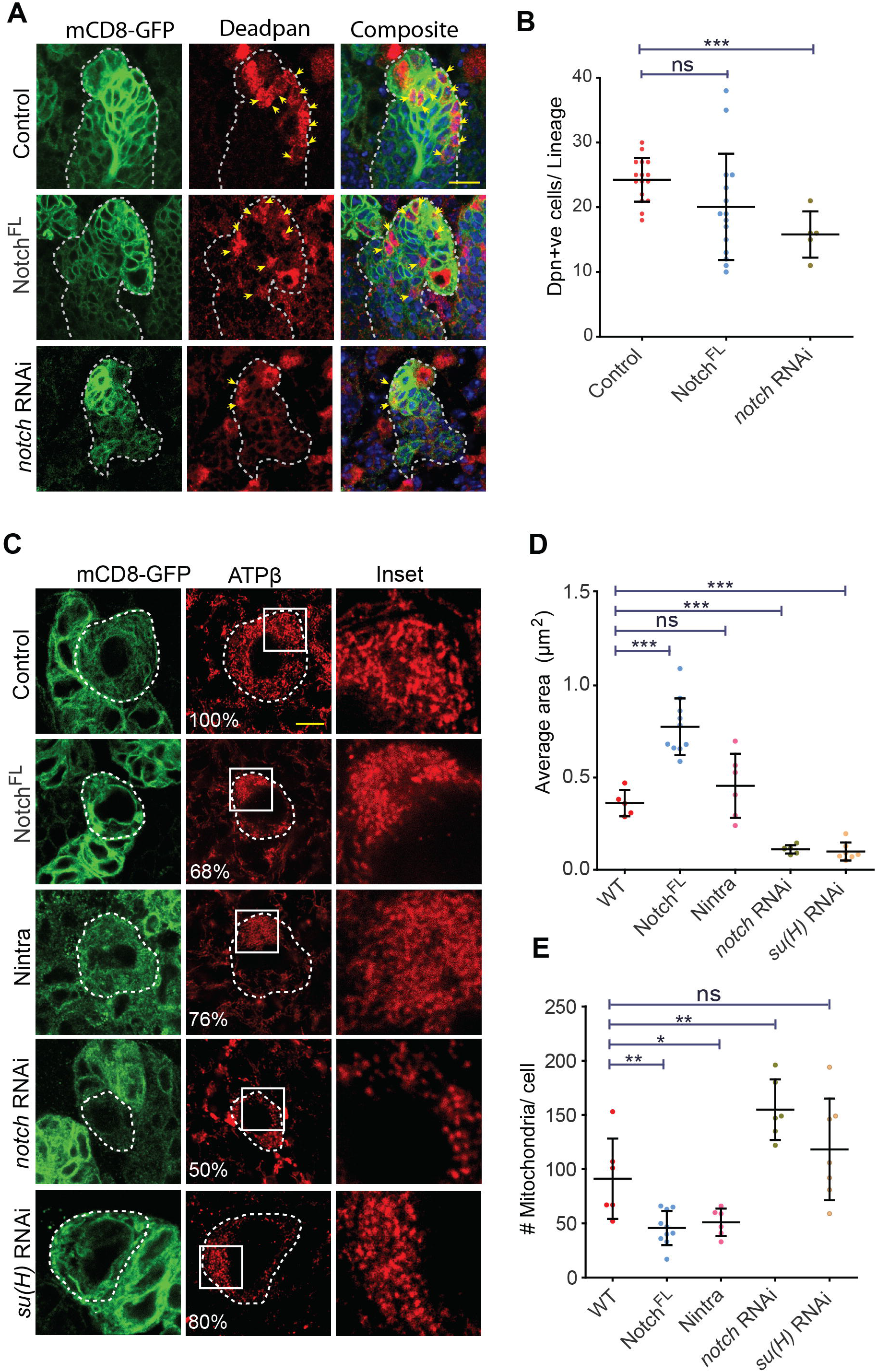
Notch regulates differentiation and fused mitochondrial morphology and in type II NBs. A-B: Type II NB lineages stained for mCD8-GFP (green) and Dpn (red, yellow arrows) show reduced mINPs in Notch downregulation (D). Analysis of mINP numbers in type II NB lineages (E) of *pnt*-Gal4, UAS-mCD8-GFP control (16 NB lineages, 4 brains), or with Notch^FL^ (Notch full length) (16,8), *notch* RNAi (5,5). Scale bar-10μm. C-E: Type II NBs (mCD8-GFP, green, yellow dotted line) showing Notch signaling mediated regulation of mitochondrial morphology (red) by ATPβ antibody using STED microscopy (A). Control (100% tubular, 75 NBs, 22 Brains), Notch^FL^ (68% clustered, 103,16), Nintra (76% clustered, 58,6), *notch* RNAi (50% fragmented, 30,14), *su(H)* RNAi (80% fragmented, 23,12). Average mitochondrial area quantification (B) in Control (6 NBs,3 brains), Notch^FL^ (10,5), Nintra (6,4), *notch* RNAi (6,4), *su(h)* RNAi (7,4). Mitochondrial number quantification (C) in control (6,3), Notch^FL^ (10,5), Nintra (6,4), *notch* RNAi (6,4), *su(H)* RNAi (7,4). Scale bar-5μm. B, D & E: Graphs show mean + sd. Statistical analysis was done by using unpaired t-test. ns-non significant, *-p<0.1, **-p<0.01, ***-p<0.001

Since Notch signaling regulated differentiation within each type II NB lineage and loss of mitochondrial fusion proteins led to loss of differentiation in the type II NB lineage, we assessed if Notch signaling regulated mitochondrial morphology in type II NBs. STED microscopy revealed thread like mitochondria evenly distributed around the nucleus in type II NBs (Figure 6C). Interestingly, overexpression of the Notch^FL^ and Nintra in type II NBs led to clustering of mitochondria on one side of the nucleus (Figure 6C). Overexpression of Notch^FL^ led to an increase in the average area of mitochondria and decrease in mitochondrial numbers per cell as compared to control NBs (Figure 6D-E). Contrarily, Notch downregulation by expression of *notch* RNAi led to an increase in fragmented mitochondria as marked by a decrease in average size of optically resolvable puncta and increase in numbers per cell. Likewise, depletion of Su(H) in type II NBs also resulted in fragmented mitochondria (Figure 6C-E). These data show that Notch activation via the canonical pathway through Su(H) leads to mitochondrial clustering and fusion whereas Notch depletion results in fragmented mitochondria.

### Mitochondrial fusion induces differentiation on Notch depletion in the type II NB lineage

Our data together shows that mitochondrial morphology is tubular in type II NBs and fusion of mitochondria alone did not give a differentiation defect but was able to suppress the differentiation defect in *opa1* mutant NBs. We further tested whether mitochondrial fusion would drive differentiation in type II NBs depleted of Notch. We combined *drp1* mutants to generate fused mitochondria along with *notch* RNAi and expressed this combination in the type II NB lineage (Figure 7A). The numbers of type II NBs remained less than controls in the *notch* RNAi, Drp1^SD^ combination and were similar to *notch* RNAi (Figure 7B). This observation further confirms that mitochondrial fusion does not affect Notch signaling in the type II NB. The loss of mINPs (Figure 7A,C) and GMCs (Figure 7D,E) was suppressed in each type II NB lineage and the numbers of mINPs and GMCs was closer to controls (Figure 7A,C,D,E). These results show that fused mitochondrial morphology can alleviate Notch signalling defects to drive differentiation in the type II NB lineage.

**Figure 7:**
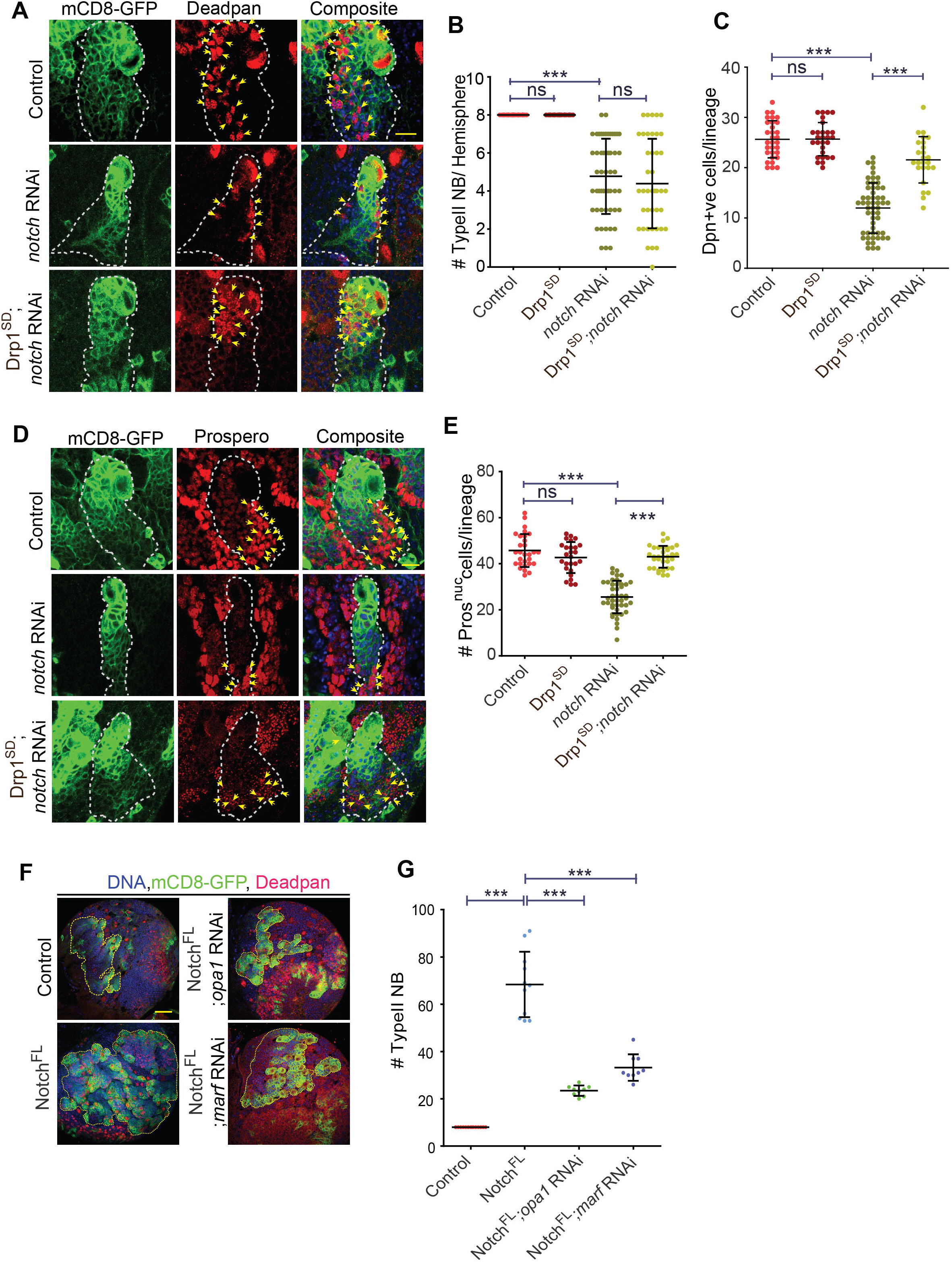
Drp1 depletion alleviates the differentiation defect seen in *notch* RNAi expressing type II NB lineages. A-C: Type II lineages (*pnt*-Gal4, mCD8-GFP, green) showing Dpn positive INPs (red, yellow arrows) (A). Quantification of number of type II NBs (B) in control (16 Brains), Drp1^SD^ (10), *notch* RNAi (9), Drp1^SD^;*notch* RNAi (5). Quantification of Dpn positive mINPs (C, yellow arrows)) in control (23 NB lineages,16 brains), Drp1^SD^ (28,10), *notch* RNAi (54,9), Drp1^SD^;*notch* RNAi (25,5). Scale bar-10μm D-E: Type II lineages (*pnt*-Gal4, mCD8-GFP, green) showing Pros positive INPs (red, yellow arrows) (D, yellow arrows). Quantification of Pros positive GMCs (E) in control (29 NB lineages,18 brains), Drp1^SD^ (26,8), *notch* RNAi (45,5), Drp1 ^SD^;*notch* RNAi (33,5). Scale bar-10μm F-G: Larval brain lobes show suppression of Notch mediated NB hyper proliferation on depletion of *opa1* and *marf* (F). Quantification of NB number (G) in *pnt*-Gal4, UAS-mCD8-GFP control (14 lobes), Notch^FL^ (11), Notch^FL^;*opa1* RNAi (9), Notch^FL^;*marf* RNAi (9). Notch^FL^ quantification is repeated from Figure 1C. Scale bar-50μm B, C, E & G: Graphs show mean + sd. Statistical analysis was done by using unpaired t-test. ns-non significant, ***-p<0.001.

To check whether fused mitochondrial morphology is essential for Notch signaling mediated NB proliferation, we depleted Opa1 and Marf along with Notch^FL^ overexpression and found that the numbers of the NBs were significantly reduced as compared to Notch^FL^ alone (Figure 7F-G). Overexpression of the intracellular domain of Notch gave rise to an increase in size of brain lobes. The brain lobe size was reduced on depletion of Opa1 and Marf on overexpression of Notch intra (Figure S5A). The NB numbers were reduced in the *Nintra; opa1* RNAi and *Nintra; marf* RNAi combination (Figure S5B-C). As expected, the clustered mitochondrial morphology seen on overexpression of Nintra (Figure 6C) was also not seen on depletion of Opa1 and Marf in the Nintra background (Figure S5D). We therefore conclude that increased Notch signaling along with fused mitochondrial architecture leads to NB proliferation.

In summary, our results show that Notch signaling increased mitochondrial fusion and mitochondrial fusion was important for differentiation in the type II NB lineage. This is also consistent with the requirement of mitochondrial fusion in NB tumors induced in *brat* and *numb* mutants, since depletion of mitochondrial fusion and not mitochondrial fission abrogates the tumor phenotype in *brat* and *numb* mutant NBs [24]. In summary our results show that mitochondrial fusion controlled by Opa1 and Marf regulates Notch signaling driven differentiation in the *Drosophila* type II NB lineage.

## Discussion

Recent evidence shows that maintenance of mitochondrial architecture is critical for cell fate determination [53–55]. Depletion of ETC components in *Drosophila* NBs leads to mitochondrial fragmentation and loss of differentiation and cancer progression [22–24]. Here we show that mitochondrial fusion proteins, Opa1 and Marf affect differentiation in the type II NB lineage (Figure 8A-B). Opa1 and Marf gave equivalent fragmentation and loss of mitochondrial activity in the type II NB. However, decreased Opa1 led to loss of both mINP and GMCs whereas decreased Marf led to loss of only GMCs in the type II lineage. Interestingly, mitochondrial outer membrane fusion ameliorated the defects in mitochondrial activity and differentiation caused by Opa1 and Marf depletion. Notch signaling regulated fused mitochondrial morphology in the type II NB and mitochondrial fusion in Notch depleted type II NB lineages led to suppression of differentiation defects. Here, we discuss our results in the following contexts: 1] the mechanisms by which Notch signaling gives rise to fused mitochondrial morphology, 2] the role of the mitochondrial morphology in mediating Notch signaling and 3] the role of mitochondrial fusion in differentiation in NBs.

**Figure 8:**
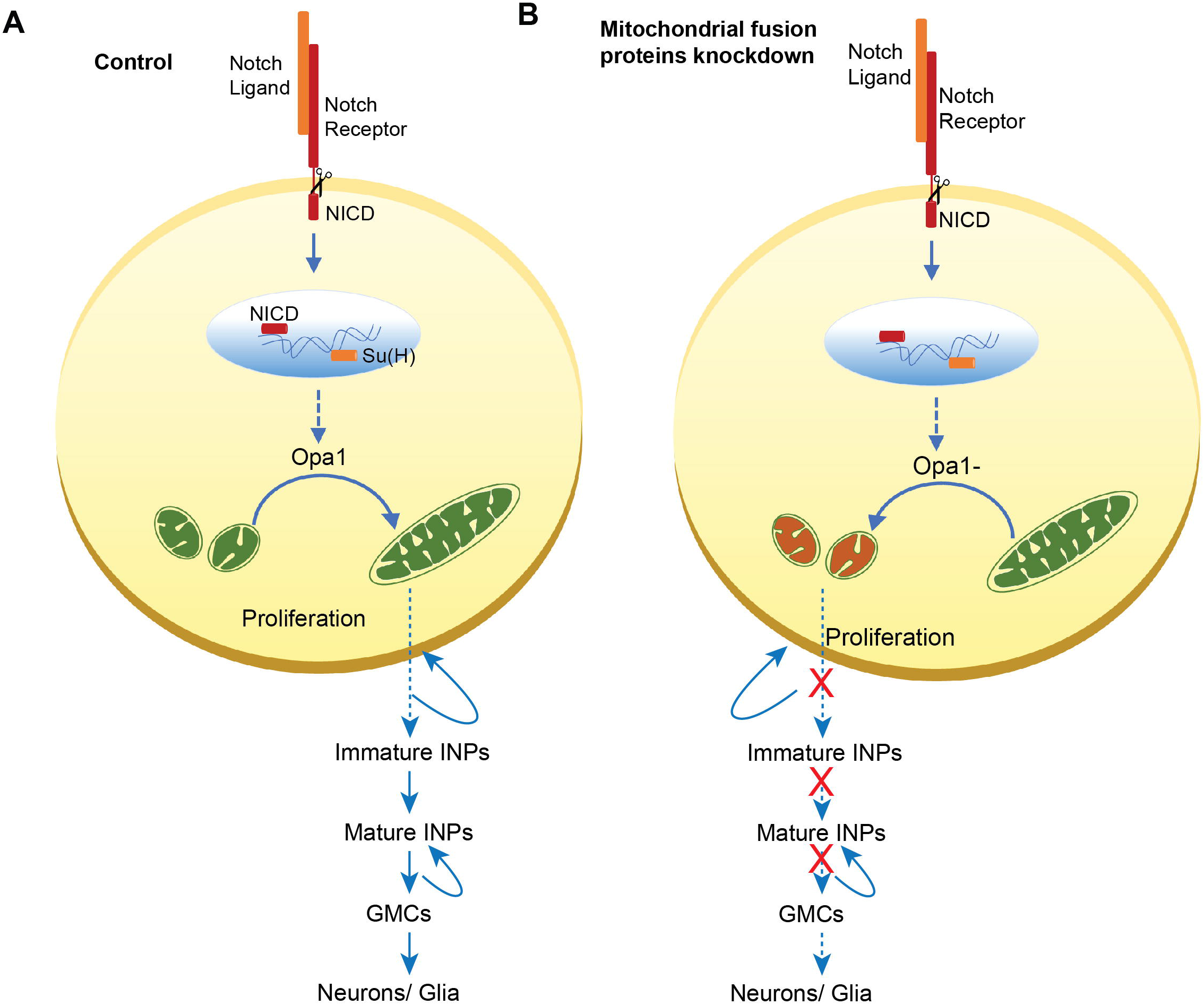
Schematic summary for regulation of type II NB differentiation by fused mitochondrial morphology. Mitochondria are maintained in a relatively tubular morphology in type II NBs in the presence of canonical Notch signaling. Mitochondrial fusion protein Opa1 is essential for NB differentiation (A). Mitochondrial fusion protein Marf is also essential for formation of GMCs. Depletion of mitochondrial fusion protein Opa1 in type II NBs leads to formation of fragmented mitochondria with lowered mitochondrial membrane potential which can potentially result in the production of defective immature INPs which further fails to differentiate into mINP and GMCs (B).

### Regulation of mitochondrial fusion by Notch signaling

At the heart of the discussion on interaction between Notch signaling and mitochondrial morphology, lies an analysis of how Notch signaling regulates mitochondrial fusion and activity. Loss of *su(H)* also showed fragmented mitochondria similar to downregulation of Notch. In addition, mitochondrial fusion alleviated the differentiation defects seen in type II NBs depleted of Notch. Since the action of both NICD and Su(H) is needed in the nucleus to activate targets downstream of Notch signaling, it is possible that Notch regulates fused mitochondrial morphology through the canonical pathway by enhancing expression of fusion genes *opa1* and *marf* or regulating activity of Opa1 and Marf by post translational modification and reducing expression or activity of Drp1. Notch signaling may also regulate mitochondrial activity by elevating components of the ETC. Recent studies show that NB tumors arising in *brat* and *numb* RNAi, which also have elevated Notch signaling, show increased transcription of Opa1 and Marf thereby causing mitochondrial fusion and increased oxidative metabolism resulting from an increase in expression of oxidative phosphorylation enzymes [24,29]. This is likely to occur due to increased Notch activity or due to the change of fate of cells in the type II NB lineage.

### Impact of mitochondrial morphology on Notch signaling

Loss of Notch signaling leads to decrease in numbers of type II NBs. Even though increased Notch signaling by *pointed*-Gal4 favored mitochondrial fusion in the Dpn positive NBs, loss of Opa1 and Marf did not affect the numbers of type I and type II NBs. This evidence suggests that mitochondrial fusion is not needed during Notch mediated formation of NBs during the embryonic stages. We found that proliferation of mINPs depended upon Notch signaling driven mitochondrial fusion. Even though the immature INP numbers in mitochondrial mutants did not change in mitochondrial fusion mutants, it is possible that mitochondrial fusion is needed to produce healthy INPs that will mature, proliferate and differentiate to form GMCs. Further, a decrease in GMCs on Opa1 and Marf depletion is possible because of defects in division of the mINPs. Loss of mitochondrial fusion led to abrogation of mitochondrial activity. How might loss of mitochondrial fusion and activity affect differentiation in the type II NB lineage? Mitochondrial morphology changes may have an effect on the fate of the cell by regulating key metabolites (Choi et al. 2020; Galloway and Yoon 2013). Mitochondrial pyruvate carrier (MPC) in the inner mitochondrial membrane is involved in transport of pyruvate from cytoplasm to the mitochondrial matrix where pyruvate undergoes oxidation via tricarboxylic acid (TCA) cycle (McCommis and Finck 2015). MPC depletion in the intestinal stem cells (ISC) results in increased ISC proliferation and loss of differentiation (Schell et al. 2017). We observed severe differentiation defects upon loss of Opa1 in NB lineage. It is possible that the defects in the inner mitochondrial membrane due to Opa1 loss affect pyruvate transport into the mitochondrial which further affect NB differentiation. It will be interesting to check whether pyruvate metabolism is affected in the mitochondrial morphology mutants.

Our preliminary analysis shows that Opa1 mutant type II NBs show a loss of Notch receptors on the plasma membrane and increase in endosomes in the type II NB and the lineage (data not shown). Numb regulates inhibition of Notch signaling in INPs [29]. It is possible that sustained activity of Numb in INPs and mINPs generated from NB proliferation leads to a decrease in Notch signaling in the mINPs. Future experiments on the defects on Notch receptor trafficking and Numb distribution in type II NBs will reveal a mechanistic link between mitochondrial morphology, activity and Notch signaling.

Notch signaling may also be affected by interaction between mitochondrial morphology and other upstream signaling pathways. Interestingly, our finding that Notch signaling maintains fused mitochondria in NBs is in striking contrast to recent literature in other systems where fused mitochondrial morphology has been correlated with loss of *notch* activity. In triple negative breast cancer (TNBC) Notch signaling enhances mitochondrial fission via *drp1* [56]. In *Drosophila* ovarian posterior follicle cells, mitochondrial fusion induces an increase in mitochondrial membrane potential and loss of Notch signaling [18,19]. Loss of *opa1* in cardiomyocytes leads to decrease in differentiation due in enhanced Notch processivity [20]. It is interesting to speculate the reasons for observing tissue specific differences in the requirement of mitochondrial fusion for Notch signaling in differentiation. We had previously found that EGFR signaling regulates fragmented morphology for appropriate Notch signaling in *Drosophila* follicle cells [18,19]. It is likely that fragmented mitochondrial morphology leads to an interaction between EGFR and Notch signaling pathways in follicle cells and possibly other cell types such as cardiomyocytes and TNBCs where Notch signaling increase occurs only on loss of mitochondrial membrane potential or fragmentation.

### Role of mitochondrial fusion in Notch driven differentiation in NBs

Mitochondrial fusion is coincident with elaborate cristae organization, increased activity and increased oxidative phosphorylation [38,57,58]. Clustered mitochondria on one side of the nucleus produced on the depletion of Drp1, did not show any defect on NB differentiation suggesting that Drp1 is dispensable for NB formation and differentiation. Interestingly, Drp1 loss driven mitochondrial hyperfusion could suppress the mitochondrial activity and differentiation defects in Opa1 and Marf mutant type II NBs. Mitochondrial fragmentation and loss of mitochondrial membrane potential is likely to cause a reduction in ATP [59]. However, ATP depletion has been reported only on combined loss of oxidative phosphorylation and glycolysis and *brat* RNAi driven tumors rely on NAD+ metabolism rather than ATP synthesis [22–24]. Since we did not see any ATP stress in Opa1 and Marf depleted type II NBs, it is interesting to speculate that other mitochondrial functions dependent upon mitochondrial activity from oxidative phosphorylation are important for type II NB differentiation. Indeed mitochondrial activity may give rise to changes in calcium buffering and key metabolites [60] thereby affecting Notch signaling in NBs and lineage cells depleted of Opa1 and Marf.

Since depletion of inner mitochondrial membrane protein Opa1 affected differentiation to a greater extent as compared to Marf, it is possible that organization of the mitochondrial ETC complexes and cristae architecture in addition to fusion are crucial for type II NB proliferation and differentiation. Organization of the cristae architecture independent of oxidative phosphorylation has been previously shown to be important for *Drosophila* germ line stem cell differentiation [61]. Fusion of the outer mitochondrial membrane may restore the inner membrane organisation and increase mitochondrial activity needed in type II NB proliferation and differentiation. Recent evidence suggests that neurodegeneration defects caused by depletion of oxidative phosphorylation in Purkinje neurons can be completely rescued by mitochondrial fusion produced by overexpression of Mfn [62]. NBs deficient of the mitochondrial ETC lead to fragmentation [23] and it will be interesting to probe if fusion of mitochondria in ETC mutants will also mitigate the differentiation defect.

In summary, we find a distinct role for mitochondrial fusion in regulation of Notch signaling in type II NB differentiation. Future studies on the mechanistic link between mitochondrial activity, change in metabolite status and Notch signaling in diverse contexts will give further insight into the interaction between Notch signaling and mitochondrial morphology in a tissue specific manner. Our studies motivate an analysis of mechanisms that regulate the interaction between mitochondrial fusion or inner membrane architecture and signaling during development and differentiation at large.

## Abbreviations

NB: neuroblast
INPs: intermediate precursor cells
GMC: ganglion mother cell
Drp1: Dynamin related protein
Opa1: Optic atrophy protein 1
Marf: Mitochondrial assembly regulatory factor

## Acknowledgements

We thank the RR lab members for discussions on this project and feedback on the manuscript. We thank the *Drosophila* facility and microscopy facility at IISER, Pune, India for support throughout this project. DD thanks IISER, Pune, India for his graduate fellowship. RR thanks DBT and IISER Pune for funding.

## Materials and methods

### Fly genetics

Fly crosses were performed in standard cornmeal agar medium and raised 29 °C. The following fly lines were used in this study: *pnt*Gal4,UAS-mCD8-GFP (Jurgen Knoblich, IMP, Vienna, Austria), *elav*>Gal4, *prospero*Gal4, *inscuteable*>Gal4, *scabrous*>Gal4, *worniu*>Gal4, *opa1* RNAi (Bloomington stock number BL32358), *opa1* RNAi (Ming Guo, UCLA), *marf* RNAi (Ming Guo, UCLA, [33]), *marf* RNAi (BL31157), *opa1* RNAi2 (BL67159), *marf* RNAi2 (BL67158, [63]), *drp*1 RNAi (BL51483), *drp1* RNAi (VDRC44155), Drp1^S193D^ (Drp1^SD^, GTPase domain mutant, acts as a dominant negative, made in the Richa Rikhy lab), *notch* RNAi (BL31383), *su(H)* RNAi (BL67928), UAS-Notch^FL^ (BL52309) and UAS-Nintra (LS Shashidhara, IISER, Pune, India), hSOD1 mutant (BL33607), mCherry RNAi (BL35785). Drp1^SD^; *opa1* RNAi, Drp1^SD^; *marf* RNAi, *notch* RNAi /CyOGFP; *opa1* RNAi /TM3SerGFP, *notch* RNAi /CyOGFP; *marf* RNAi/TM3SerGFP lines were generated using standard genetic crosses. *worniu*>Gal4,UAS-mCD8-GFP (*wor*-Gal4) was used to express transgenes in all the larval NBs in the third instar larval brain and *pointed*-Gal4 (*pnt*-Gal4) was used to express transgenes in the type II NBs in the larval brain.

### Immunostaining of larval brain

Wandering third instar larvae were dissected in Schneider's medium and immediately fixed in 4% PFA solution for 25 minutes at room temperature (RT). The brains were washed subsequently with 1X PBS with 0.1% Triton X-100 (PBST) for 30 minutes at RT. They were blocked with 1% BSA for 1hr at RT. The brains were stained with the appropriate primary antibody overnight at 4°C. They were then washed 3 times with 0.1% PBST (first wash for 20 minutes and remaining for 10 minutes each). An appropriate fluorescently coupled secondary antibody was added for 1hr at RT followed by three washes with 0.1% PBST (first wash for 20 min and remaining for 10 min each) and mounted in Slow-Fade Gold (Molecular Probes).

The following dilutions were used for the primary antibodies: chicken anti-GFP (1:1000, Invitrogen), anti- ATPβ (1:200, Abcam), anti- Deadpan (1:150, Abcam), anti-Prospero (1:25, DSHB), anti- Miranda (1:600, Abcam), anti-Cytochrome C (1:200, Cell Signaling), anti-Elav (1:100, DSHB), anti-cleaved Caspase3 (1:100, Cell Signaling), anti-phosphohistone 3 (1:100, Invitrogen), anti-phosphoAMPK (1:200, Invitrogen). Hoechst (1:1000, Molecular Probes) was used to label DNA. Fluorescently coupled secondary antibodies (Molecular Probes): anti-Chicken 488, anti-Rat 568/633/647, anti-Rabbit 568, anti-Mouse 568/633 were used in 1:1000 dilution.

### DHE uptake for live imaging of ROS

Dissected third instar larval brains were treated with Dihydroethidium (DHE) (1:1000, Molecular Probes) in Schneider's medium for 15 minutes at RT and then washed with Schneider's medium for 10 min. Brains were mounted in LabTek chambers containing Schneider's medium and imaged immediately using Zeiss LSM 710 with a 63x/1.4NA oil objective using a DPSS (561 nm) laser and dihydroethidium-1 filter settings in the Zeiss2010 software. The laser power, acquisition speed, frame size and gain were kept the same for both control and mutant. The laser power and gain were adjusted to keep the range of acquisition between 0-255 on an 8-BIT scale.

### 2-Deoxy glucose treatment

Third instar larval brains were treated with 500μM 2-DG in Schneider’s medium for 1hr. Control and treated brains were processed for pAMPK immunostaining as mentioned above.

### Mitochondrial membrane potential estimation with TMRM in live brains

Third instar larval brains were dissected in Schneider’s medium. Then they were treated with Tetramethylrhodamine, methyl ester (TMRM) (100nM, Thermo fisher Scientific) for 30 mins at room temperature. Treated brains were mounted in a LabTek chamber containing Schneider's medium and imaged immediately using Zeiss LSM710 with a 63x/1.4NA oil objective. A DPSS (561 nm) laser and RFP/TRITC filter was used for the detection of TMRM signal.

### TUNEL assay for detection of apoptotic cells

Third instar wandering larvae were dissected in Schneider's medium and fixed in 4% PFA for 25 mins followed by washing with 0.1% PBST for 20 mins at RT. Brains are then washed twice with 1XPBS for 2 min. The terminal transferase (TdT) reaction was performed as follows: First brains were treated with TdT reaction buffer and incubated for 10min followed by treatment of freshly made TdT reaction buffer cocktail (TdT reaction buffer, 5-Ethynyl-dUTP, TdT) for 60 min at 37°C and at 500rpm. After the TdT reaction, brains were washed twice with 3% BSA at RT for 5 min. Then Click-iT reaction was performed by adding click iT reaction cocktail (Click-iT reaction buffer, Click-iT reaction buffer additive) for 30min at RT. At this step samples were protected from light. Brains were washed with 3% BSA twice for 5 min after removing Click-iT reaction buffer cocktail and then incubated with Hoescht (1:1000) for DNA staining followed by washing with 1XPBS for 10 min. Samples mounted in Slow-Fade Gold (Molecular probes) and subsequently imaged using Zeiss LSM710 with a 40x/1.4NA oil objective and DPSS laser (561nm). Induction of apoptosis by UAS-Hid line driven by *pointed* Gal4 was used as positive control. TUNEL positive nuclei were counted and plotted using GraphPad Prism 5 software.

### Microscopy and Image acquisition

#### Imaging of fixed samples using a confocal microscope

Confocal microscopy of fixed samples was done at room temperature using LSM710 or LSM780 inverted microscope (Carl Zeiss, Inc. and IISER Pune microscopy facility) with a Plan apochromat 40x 1.4NA and 63x 1.4NA oil objective. Images were acquired using the Zen2010 software at 1024×1024 pixels with an averaging of 4 and acquisition speed 7. Fluorescence intensity was kept within 255 on an 8-bit scale. Following lasers were used for the excitation of different fluorophores during fixed sample imaging: Diode laser for Hoescht, Argon laser line at 488nm for Alexa Fluor 488, DPSS laser for Alexa Fluor 568, HeNe (633nm) for Alexa Fluor 633 and Alexa Fluor 647. The representative image for each type II lineage in the figures is shown from the center of the lineage and comprises the maximum number of cells in the lineage.

#### Stimulated Emission-Depletion (STED) microscopy for imaging mitochondrial morphology

Super-resolution microscopy was done for visualizing mitochondrial morphology within type II NBs using the Leica TCS SP8 STED 3X Nanoscope with a 100x/1.4NA oil objective. Images were acquired using the LasX software at 1024×1024 pixels to keep pixel size 20-25nm, an averaging of 4, an acquisition speed of 200 and a zoom of 4.5. The Alexa Fluor 488 and 568 were excited with Argon 488 nm and Diode 561 nm lasers respectively and emission were collected with hybrid detectors for GFP and mitochondria labeled with ATPβ antibody respectively. The 561 nm excitation laser with the 775nm depletion laser was used for stimulated emission-depletion for visualizing mitochondria by super resolution. Fluorescence intensity was kept within 255 on an 8-bit scale using the LUT mode to avoid over-saturated pixels.

### Image analysis and statistics

#### Neuroblasts number analysis

NBs in a single hemisphere were counted by using Miranda as a NB marker across different mitochondrial dynamics mutants and compared with controls. Type II NBs were counted by using mCD8-GFP positive lineages expressed under *pointed* Gal4. Non-parametric student t-test was performed for statistical analysis.

#### ROS intensity analysis

Fluorescence intensity of DHE uptake in control and mutant type II NB (mCD8-GFP positive) along with their neighboring NB (mCD8-GFP positive) were quantified by drawing a region of interest using a free hand tool in ImageJ. Ratios of DHE fluorescence intensity of GFP positive NB to a neighboring GFP negative NB present in the same optical plane were plotted using GraphPad Prism software. Non-parametric student t-test was performed for statistical analysis

#### pH3 Analysis

pH3 positive cells were counted in each type II NB lineage for control and mutant brains and plotted using GraphPad Prism software. Non-parametric student t-test was performed for statistical analysis.

#### Deadpan and Prospero quantification for mINPs and GMCs

Immature INPs were identified and quantified as Dpn-and Pros-cells in each type II NB lineage labelled with mCD8-GFP and Dpn and Pros. Dpn positive mINPs were counted in all sections in each type II NB lineage in control and mutant brains. Numbers of mINPs were plotted using GraphPad Prism software and non-parametric student-t-test was performed for statistical analysis. Nuclear Pros containing GMCs were counted in the type II NB lineage. For Pros analysis in each brain hemisphere for larval NBs of the type I and type II lineage, the maximum intensity optical plane was selected from anterior and posterior regions. We obtained the average area for each Pros positive nucleus from 5 GMCs in each sample. The total Pros positive area was determined by using the threshold tool in ImageJ. To extract the number of Pros positive GMCs, we divided the total area by average area of a single GMC.

### TMRM analysis

TMRM intensities were computed in the entire type II NB in control, *opa1* RNAi and *marf* RNAi expressing brains and in the clustered mitochondria in Drp1^SD^, Drp1^SD^;*opa1* RNAi and Drp1^SD^;*marf* RNAi expressing brains using ImageJ. Average intensity was computed from the GFP positive type II NB and expressed as a ratio to the intensity seen in the neighboring control GFP negative NB. Relative intensities were then plotted using GraphPad Prism software.

### pAMPK analysis

Average pAMPK intensities in control and mutant GFP positive type II NB and neighboring GFP negative NBs were measured using ImageJ software. pAMPK fluorescence was plotted as a ratio to the neighboring control cells by using GraphPad Prism software. Larval brain treatment of 2-DG showed an overall increase in antibody fluorescence and the intensity in the GFP positive type II NB was used as positive control to induction of pAMPK.

### Mitochondrial number and area analysis

Control and mutant brains were immunostained with ATPβ antibody. A qualitative observation of mitochondrial distribution as tubular, clustered and dispersed was made by visually observing NBs in various genotypes. Quantitative measurements for size and number of mitochondria were also done by using thresholding tool in imageJ software. A single optical plane was selected approximately in the middle of the type II NB (section with the highest diameter) where the nucleus was prominently visible and the density of the mitochondria was high. All mitochondrial particles which were above size cut off of 0.12 μm^2^ were analyzed for size. To clearly resolve mitochondrial particles in different genotypes we used a watershading tool from ImageJ.

### TUNEL analysis

TUNEL positive nuclei per central brain region or type II NB lineage were counted in lobes of control and *opa1* RNAi and plotted using GraphPad Prism software. We induced apoptosis by expressing UAS-Hid with *pnt* Gal4 as a positive control for TUNEL assay.

## Supplementary figure legends

**Figure S1:**
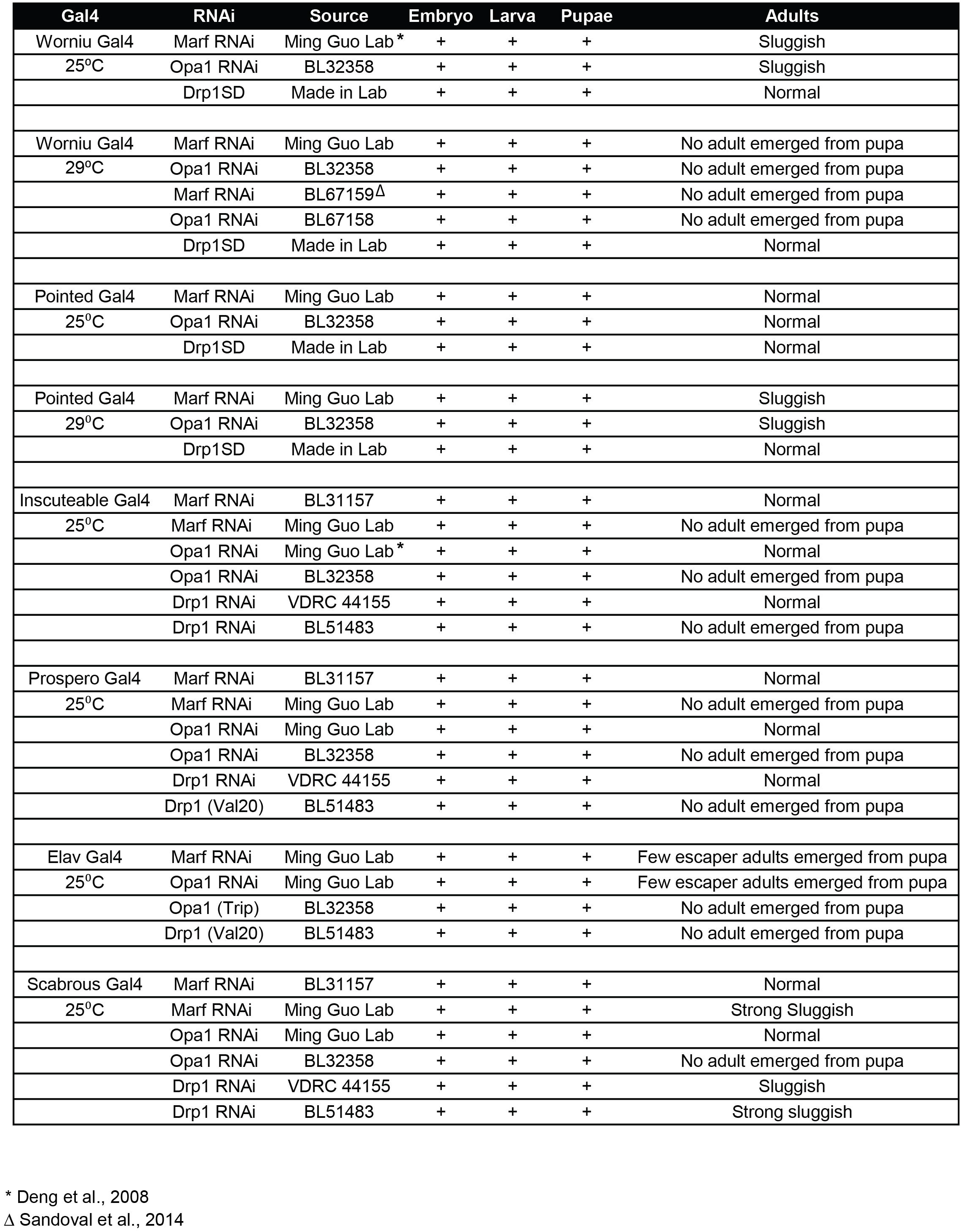
Table showing analysis of mitochondrial morphology protein knockdown with different neuronal Gal4s. Various Gal4 drivers were crossed with mutants of mitochondrial morphology genes at 25 or 29 °C and lethality or behavioral phenotype was recorded in the adult. *worniu*-Gal4 (*wor*-Gal4), *inscuteable*-Gal4, *prospero*-Gal4 and *scabrous*-Gal4 expresses the Gal4 in all NBs, *pointed*-Gal4 (*pnt*-Gal4) expresses in type II NBs and *elav*-Gal4 expresses in neurons. Adult flies from crosses with *wor*-Gal4 (25 °C) and *pnt*-Gal4 (29 °C) with *opa1* RNAi and *marf* RNAi were sluggish and the numbers obtained were at the expected frequency, no lethality was seen at the pupal stage. *elav*-Gal4 crosses gave lethality and few adults emerged. The *opa1* RNAi BL32358 and *opa1* RNAi2 BL67158 BL gave stronger phenotypes as compared to *opa1* RNAi Ming Guo lab with *inscuteable*-Gal4, *prospero*-Gal4, *elav*-Gal4 and *scabrous*-Gal4. The *marf* RNAi Ming Guo lab and *marf* RNAi2 BL67159 gave stronger phenotypes as compared to marf RNAi BL31157 with *inscuteable*-gal4, *prospero*-Gal4 and *elav*-Gal4. We chose *opa1* RNAi BL32358, *opa1* RNAi2 BL67158, *marf* RNAi Ming Guo lab and *marf* RNAi BL67159 for further analysis. The Drp1 RNAi from VDRC did not show phenotypes and the Drp1 (BL51483) gave inconsistent results, hence we used a lab generated GTPase domain mutant Drp1^SD^ for further analysis.

**Figure S2:**
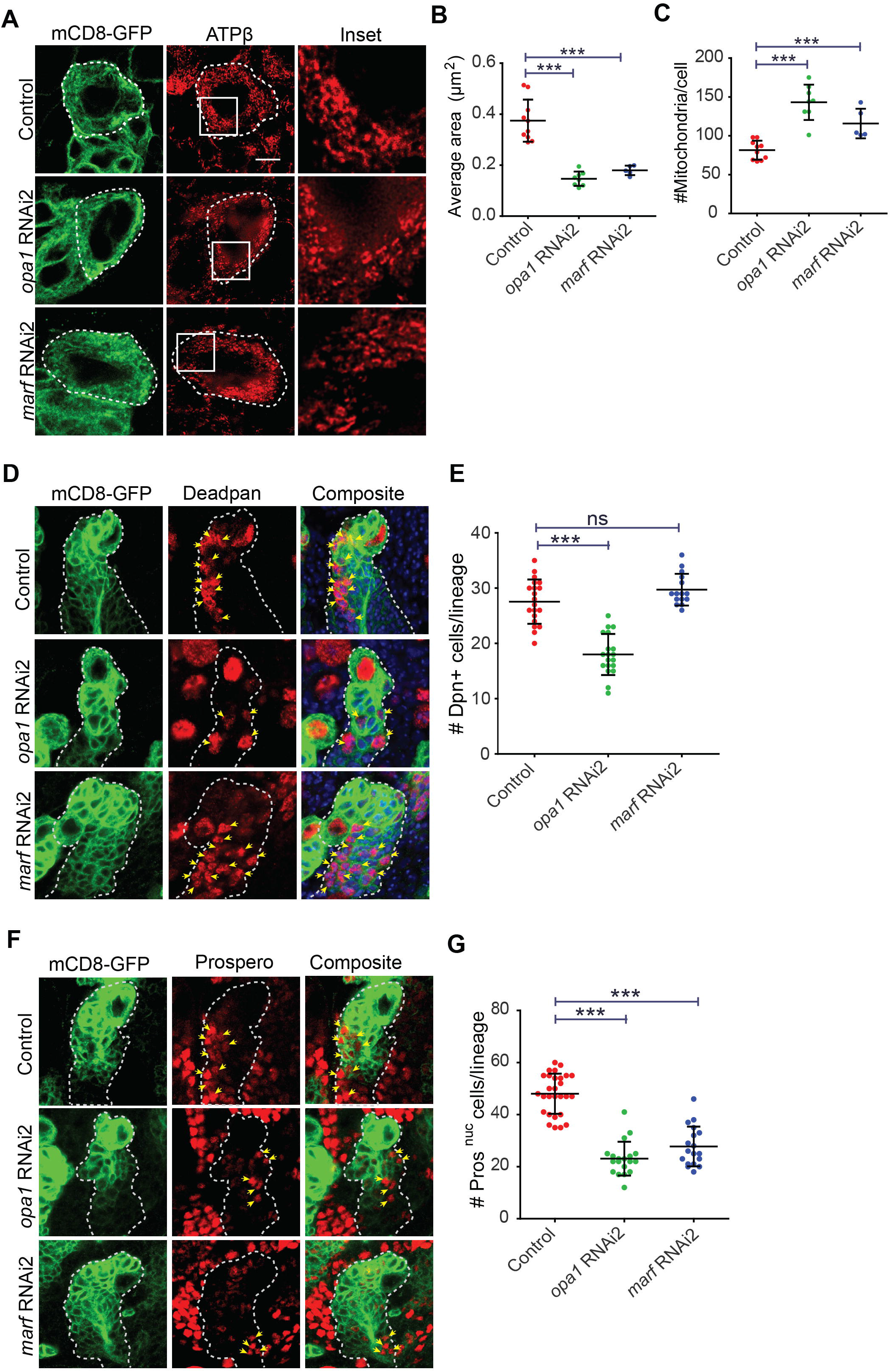
Depletion of mitochondrial fusion proteins Opa1 and Marf result in mitochondrial fragmentation and loss of type II NB differentiation. A-C:Mitochondrial morphology and distribution in type II NBs (white dotted line, magnified area shown in the panel on the right) stained with ATPβ (red) antibody using STED super resolution microscopy is shown in representative images with zoomed inset in the right panel (A). Control (100% tubular, 75 NBs, 22 brains), *opa1* RNAi2 (100% fragmented, 14,4), *marf* RNAi2 (100% fragmented, 14,4). Average mitochondrial area quantification from type II NBs (B) in control (10 type II NBs, 4 brains), *opa1* RNAi2 (8,4), *marf* RNAi2 (5,3). Mitochondrial number quantification in control (10,4), *opa1* RNAi2 (8,4), *marf* RNAi2 (5,3) (C). Scale bar-5μm D-E:Type II NB lineages (yellow dotted line) showing expression of mCD8-GFP (green) and Dpn (red, yellow arrows) (D). Quantification of Dpn positive mINPs (E) in control (20 NB lineages, 5 brains), *opa1* RNAi2 (19,4), *marf* RNAi2 (15,4). Scale bar-10μm. F-G:Type II lineages (*pnt*-Gal4, mCD8-GFP, green) showing Pros positive INPs (red, yellow arrows) (D, yellow arrows). Quantification of Pros positive GMCs (E) in control (28 NB lineages,8 brains), *opa1* RNAi2 (19,4), *marf* RNAi2 (18,4). Scale bar-10μm. B-C, E, G: Graphs show mean + sd. Statistical analysis is done using an unpaired t-test. ns-non significant, **-p<0.01, ***-p<0.001.

**Figure S3:**
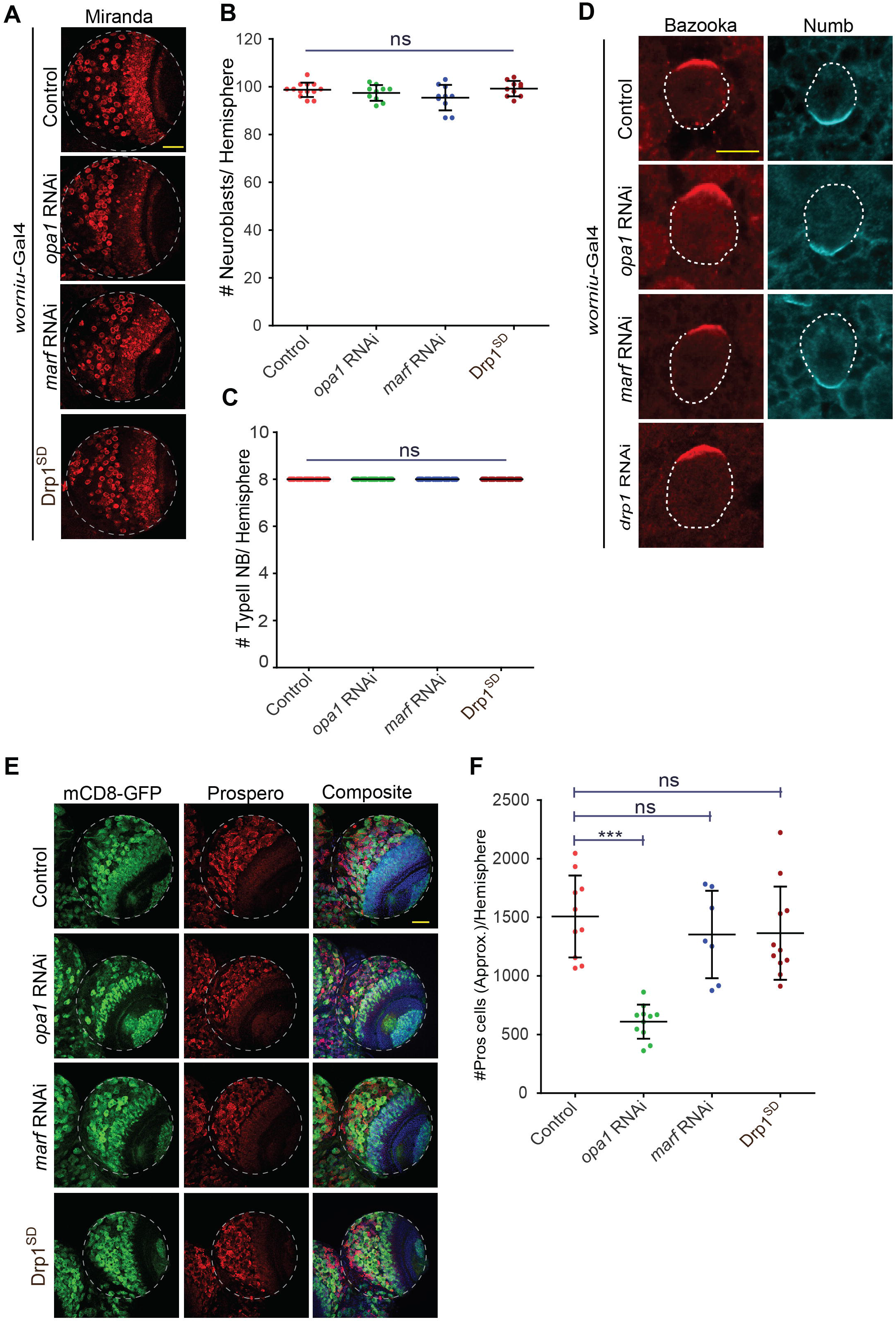
NB polarity and number are unaffected by knockdown of mitochondrial morphology proteins. A-B: Representative confocal images of larval brain lobes stained for Miranda (red) show no change in NB number (A). Expression of *opa1* RNAi, *marf* RNAi and Drp1^SD^ mutant were done by *wor*-Gal4. Quantification of NB number in larval brain lobes (B) of control (13 lobes,13 brains), *opa1* RNAi (10,10), *marf* RNAi (10,10), Drp1^SD^ (10,10). Scale bar-50μm C: Number of type II NBs and their progenies per lineage in control, *opa1* RNAi, *marf* RNAi, Drp1^SD^ (n= 30 type II NB lineages,15 brains each). D: Representative images showing apical and basal localization of Bazooka and Numb respectively in control and mitochondrial dynamics mutants. Scale bar-10μm E-F: Representative images of brain lobes showing reduced Pros positive GMC population in *wor*-Gal4 *opa1* RNAi (E). Analysis of Pros positive GMCs (F) in control (10 lobes,10 brains), *opa1* RNAi (11,11), *marf* RNAi (7,7), Drp1^SD^ (11,11).Scale bar-50μm B, C & F: Graphs show mean + sd. Statistical analysis was done by using unpaired t-test. ns-non significant, ***-p<0.001

**Figure S4:**
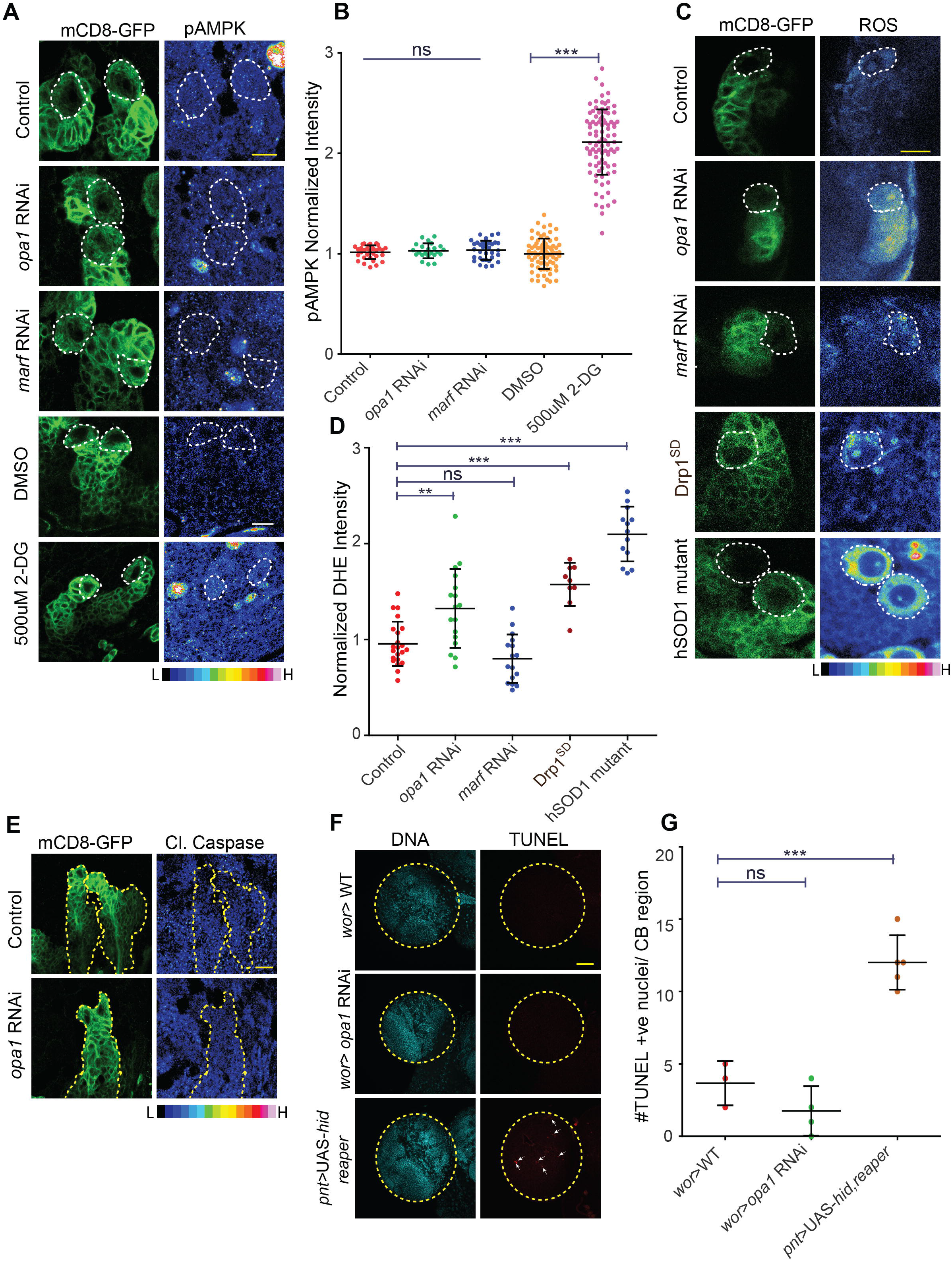
Depletion of Opa1 does not cause ATP stress and apoptosis in type II NB lineages. A-B: Representative images of type II NBs expressing mitochondrial fusion mutants along with *pnt*-Gal4, UAS-mCD8-GFP did not show change in levels of pAMPK (pAMPK fluorescence is shown as rainbow scale) (A). Controls, mutants and 2-DG treated brains were imaged for pAMPK fluorescence at the same time under the same imaging conditions. Type II NBs are marked by white dotted lines while lineages are marked by expression of mCD8-GFP (green). Analysis of pAMPK intensity normalised with neighboring control cells (B). Control (41 NBs,14 brains), *opa1* RNAi (33,8), *marf* RNAi (23,8), DMSO (76,8), 2-DG (91,9). C-D: Representative images showing increased levels of ROS (rainbow scale) in type II NB (position marked by white dotted line) using *pnt*-Gal4, UAS-mCD8-GFP with mitochondrial dynamics mutants (C). Analysis of relative DHE fluorescence as a ratio to neighboring cells (D) in type II NBs, control (22 NBs,6 Brains), *opa1* RNAi (17,8), *marf* RNAi (17,8), Drp1^SD^ (9,5), hSOD1 mutant (13,3). E-G: Representative images of type II NB lineages (*pnt*-Gal4, UAS-mCD8-GFP, green) stained for cleaved caspase and shown in heatmap in *opa1* RNAi (E) control (10 NB lineages,6 Brains), *opa1* RNAi (9,6). Fluorescence confocal images of brain hemispheres showing no significant change in TUNEL positive nuclei (red) (F) in *opa1* RNAi expressed in all NBs with *wor*-Gal4. Expression of UAS-hid shows a significant increase in TUNEL positive cells when expressed with *pnt*-Gal4, mCD8-GFP in the type II NB lineage. Quantification of TUNEL positive nuclei (G) in *wor*>WT(3 brains), *wor*> *opa1* RNAi (4), *pnt*> UAS-*hid reaper* (5). Statistical analysis was performed using unpaired t-test. ns- non significant, ***-p<0.0001 B, D & G: Graphs show mean + sd. Statistical analysis was done by using unpaired t-test. ns-non significant,**-p<0.01 ***-p<0.001

**Figure S5:**
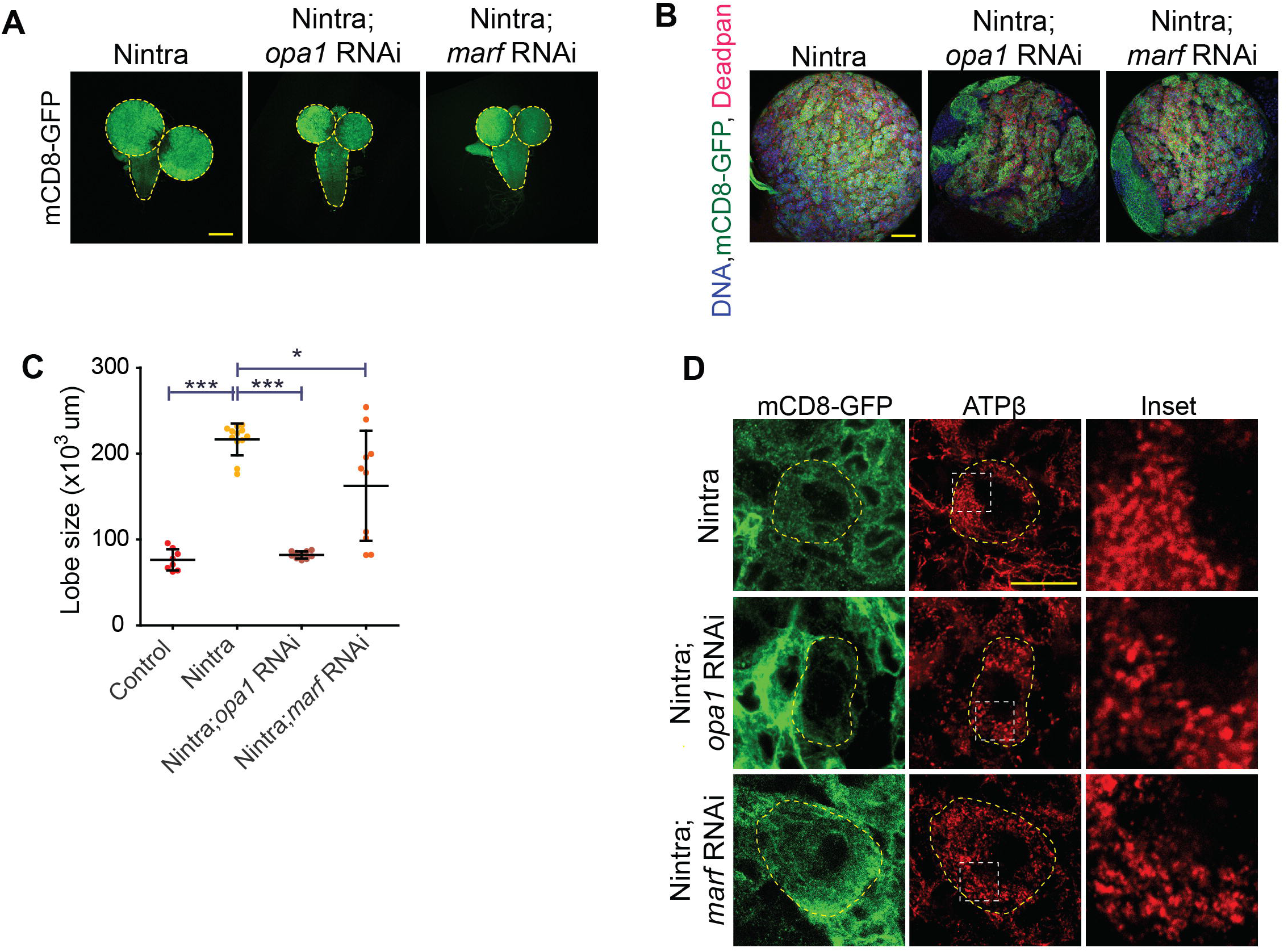
Nintra overexpression driven NB proliferation is partially reversed by additional depletion of Opa1 or Marf. A-C: Larval brain lobes (CD8-GFP, green) (A) increase in size on Nintra overexpression which is rescued by *opa1* RNAi and *marf* RNAi expression. Representative images of larval brain hemisphere containing mCD8-GFP (green, A, B) DNA (blue, B) and Dpn (red, B) showing partial rescue of NB hyperproliferation in Nintra expression by *opa1* RNAi and *marf* RNAi (B). Nintra (8 lobes, 8 brains), Nintra;*opa1* RNAi (10,10), Nintra; *marf* RNAi (10,10). Scale bar-50μm. Quantification of lobe size (C) in control (8 brains), Nintra (12), Nintra;*opa1* RNAi (10), Nintra; *marf* RNAi (12). C: Graph shows mean + sd. Statistical analysis was performed using unpaired t-test. *-p<0.05, ***-p<0.001. Scale bar-200μm. D: Representative superresolution STED images of mitochondria showing clustered morphology in Nintra mutant expressed with *pnt*-Gal4, UAS-mCD8-GFP and punctate appearance in *opa1* and *marf* RNAi expressing type II NBs in the Nintra background. Nintra (76% tubular, 58 Neuroblasts, 6 Brains), Nintra;*opa1* RNAi (100%, 95, 20), Nintra*;marf* RNAi (100%, 98, 14). Scale bar-10μm

